# Bud13 Promotes a Type I Interferon Response by Countering Intron Retention in Irf7

**DOI:** 10.1101/443820

**Authors:** Luke S. Frankiw, Devdoot Majumdar, Christian Burns, Annie Moradian, Michael J. Sweredoski, David Baltimore

## Abstract

Intron retention (IR) has emerged as an important mechanism of gene expression control. Despite this, the factors that control IR events remain poorly understood. We observed consistent IR in one intron of the Irf7 gene and identified Bud13 as an RNA-binding protein that acts at this intron to increase the amount of successful splicing. Deficiency in Bud13 led to increased IR, decreased mature Irf7 transcript and protein levels, and consequently to a dampened type I interferon response. This impairment of Irf7 production in Bud13-deficient cells compromised their ability to withstand VSV infection. Global analysis of Bud13 knockdown and BUD13 cross-linking to RNA revealed a subset of introns that share many characteristics with the one found in Irf7 and are spliced in a Bud13-dependent manner. Deficiency of Bud13 led to decreased mature transcript from genes containing such introns. Thus, by acting as an antagonist to IR, Bud13 facilitates the expression of genes at which IR occurs.

## INTRODUCTION

Three forms of alternative processing of a pre-mRNA have been described: differential inclusion of an exon, alternative splice site selection, and intron retention (IR). The latter, IR, has emerged as a previously underappreciated mechanism of post-transcriptional gene regulation. Unlike the two alternative splicing events, IR rarely contributes to proteomic diversity.^1^ However, IR events have the ability to act as negative regulators of gene expression by: (1) delaying onset of gene expression by slowing down splicing kinetics, (2) increasing potential nuclear degradation by nuclear exosomes, (3) increasing potential cytoplasmic degradation by nonsense mediated decay.^2^

Recent genomic studies suggest IR plays an important role in the regulation of gene expression in a wide range of processes including cellular differentiation^3^,^4^ and tumorigenesis.^5^ Further, widespread IR throughout mouse and human cell and tissue types has led to the idea that IR events act to functionally “tune” the transcriptome of a cell.^6^ However, with few exceptions, the factors that control IR events and thus potentially shape gene expression programs of cells, remain poorly understood.

Irf7 is an interferon-inducible master regulator of the type-I interferon-dependent immune response and is crucial to the production of interferon-α and β.^7^ Aberrant Irf7 production is linked to a wide range of pathologies, from life-threatening influenza^8^ to autoimmunity^9^, because precise regulation of Irf7 ensures a proper immune response. Notably, intron 4 of Irf7 is short, GC-rich, and has a poor splice donor sequence, characteristics shared by many poorly spliced introns. We and others have previously shown that intron 4 of Irf7 splices inefficiently^10^, affecting gene expression and opening a new line of inquiry as to the mechanism of IR regulation in *Irf7.*

Using RNA antisense purification-mass spectrometry (RAP-MS)^11^, we identified the protein Bud13 as one that regulates IR in Irf7. Bud13 was found to aid splicing efficiency and expression of the Irf7 mature transcript and protein, thus increasing the downstream type-I interferon-dependent immune response. We show that Irf7 is able to trigger a robust interferon response in the presence but not in the absence of Bud13. Further, Bud13 was found to increase the splicing efficiency of a multitude of other junctions with similar characteristics to the one found in Irf7. By aiding in splicing efficiency, Bud13 limits intron retention and increases gene expression levels of transcripts containing Bud13 dependent junctions.

## RESULTS

### Irf7 contains an intron that splices poorly following stimulation

To study the role of mRNA splicing during an innate immune response, we sequenced the RNA from mouse bone marrow-derived macrophages (BMDMs) stimulated with either TNFα, IFNα, or Poly(I:C). From this sequencing, we identified an increased number of intronic reads in the fourth intron of the most abundant transcript of Irf7 independent of time or stimulant (Fig. 1A, I, J). A variety of features of this intron make it a likely candidate for retention.^6^ It is extremely small at 69 nucleotides and has a high G/C content in both the flanking exons and within the intron itself. Furthermore, the intron contains a ‘weak’ 5’ splice site, one that deviates significantly from a consensus splice site sequence. This is quantified using a maximum entropy model to calculate the splice site quality score (Fig. 1F).^12^

**Figure 1:**
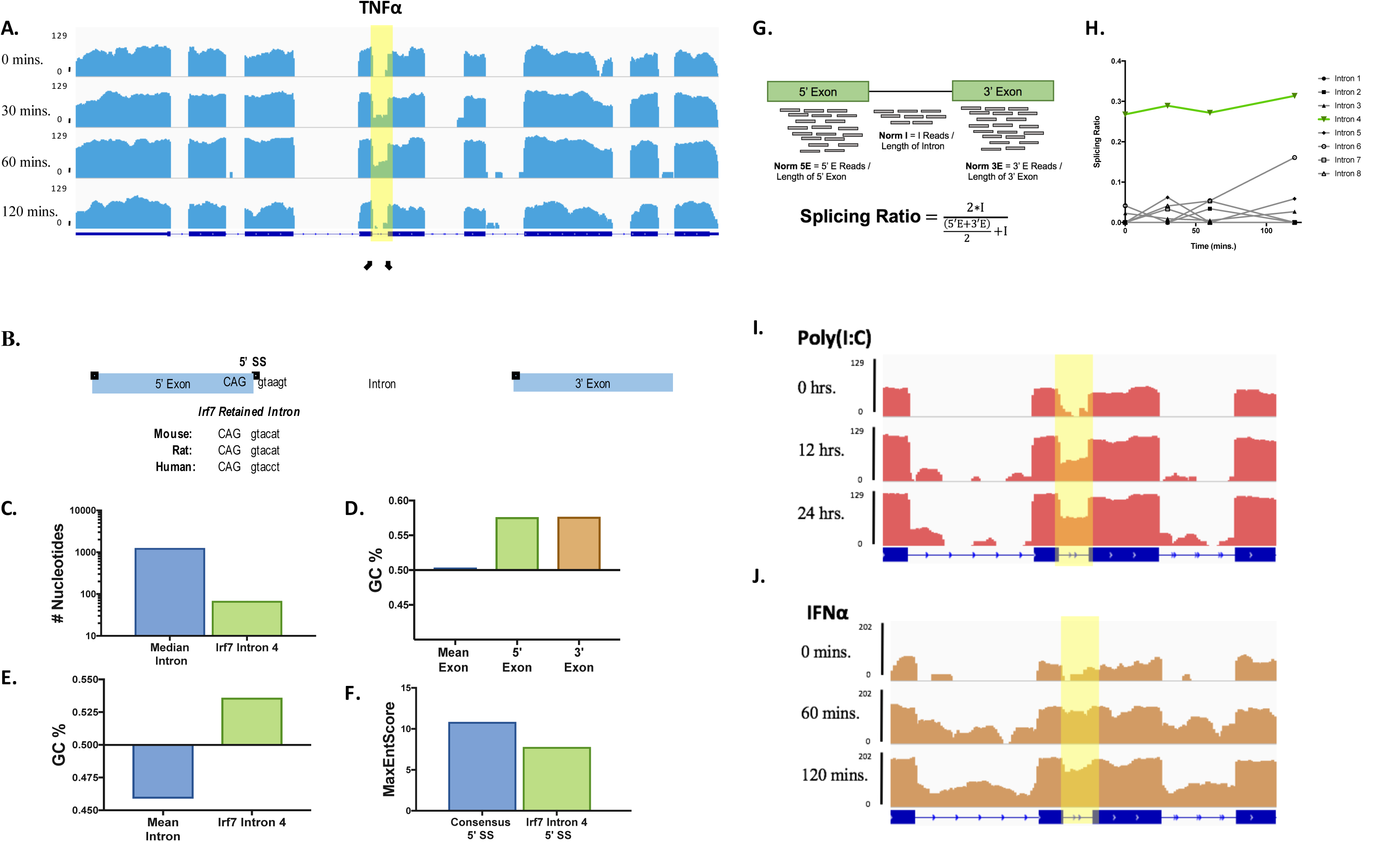
*Irf7* contains a weak intron that is following many forms of stimulation. **(A)** Histogram of mapped reads corresponding to the TNFα-induced expression of Irf7. The poorly spliced fourth intron is highlighted. For all read density plots, reads are histogrammed in log10 scale and normalized to the maximum value across the stimulation. **(B)** Comparison of Irf7 splice donor and acceptor sites in mice, rats, and humans. **(C)** Comparison of median intron length across all annotated transcripts to the length of Irf7 intron 4. **(D)** Comparison of mean exon G/C% across all annotated exons compared to the G/C% of the exons directly upstream and downstream of Irf7 intron 4. **(E)** Mean intron G/C% across all annotated introns as compared to Irf7 intron 4. **(F)** Comparison of donor splice site strength (calculated using a maximum entropy model; see methods) for the consensus donor splice site as compared to the Irf7 donor splice site. **(G)** Outline of Splicing Ratio (SR) metric. **(H)** Splicing ratio for all introns in Irf7 plotted against time stimulated with TNFα. **(I, J)** Histogram of mapped reads corresponding to the IFNα **(I)** and poly(I:C) **(J)** induced expression of Irf7 focused on the slow splicing fourth intron.

To quantify the extent of retention across RNA-seq data-sets, we use a metric we designate the “splicing ratio” (SR) (Fig. 1G; see methods), which is a length normalized ratio of intronic reads to total reads at each junction. Low SR values indicate a junction is primarily spliced whereas high SR values indicate a junction is primarily unspliced. Using this metric, we quantified the extent of retention for all junctions in the most abundant Irf7 transcript. We observed that for all types of stimulation, the retention of the fourth intron of the transcript is much greater than that seen for any of the other introns (Fig. 1H and S1 A, B). This intron remains poorly spliced despite the fact that there is clear excision of neighboring introns. It is worth noting that quantitation of the IFNα stimulation shows increased intronic signal throughout the Irf7 transcript. This increased intronic signal is due to faster and stronger induction of Irf7 via stimulation with IFNα and as such, an increase in the amount of pre-mRNA at a given stimulation time-point. Despite this increase in intronic signal throughout the transcript, we observed a corresponding increase in the level of retention for the poorly spliced fourth intron (Fig. 1J, S1B). Thus, we conclude this intron of Irf7 splices poorly following many forms of stimulation.

### RAP-MS identifies Bud13 as an RNA binding protein that interacts with IRF7 mRNA

To understand how cells handle a retained intron, we sought to identify trans-acting proteins that might affect the process using RNA Antisense Purification followed by Mass Spectrometry (RAP-MS) (Fig. 2A)^11^. RAP-MS employs antisense biotin-containing ssDNAs complementary to *Irf7* exons to purify the proteins associated with the total pool of *Irf7* transcripts, containing both nascent pre-mRNAs and mature mRNA. Using this proteomic approach, we identified the RNA-binding protein Bud13 to be highly enriched (~6-fold) on Irf7 transcripts as compared to β-*actin* transcripts, which were used as a control (Fig. 2B). Bud13 has been characterized in yeast as a member of the Retention and Splicing complex (RES),^13^ forming a trimeric complex with Pml1p and Snu17p, and aids in the splicing and nuclear retention of a subset of transcripts. It is not well characterized in mammalian systems. We captured a variety of other known RNA-binding proteins (Pum2, Prpf40a, Son); however, no other protein was enriched greater than two fold on Irf7 transcripts. We observed specificity in the RNA antisense purification for the intended transcripts (Fig. 2C).

**Figure 2:**
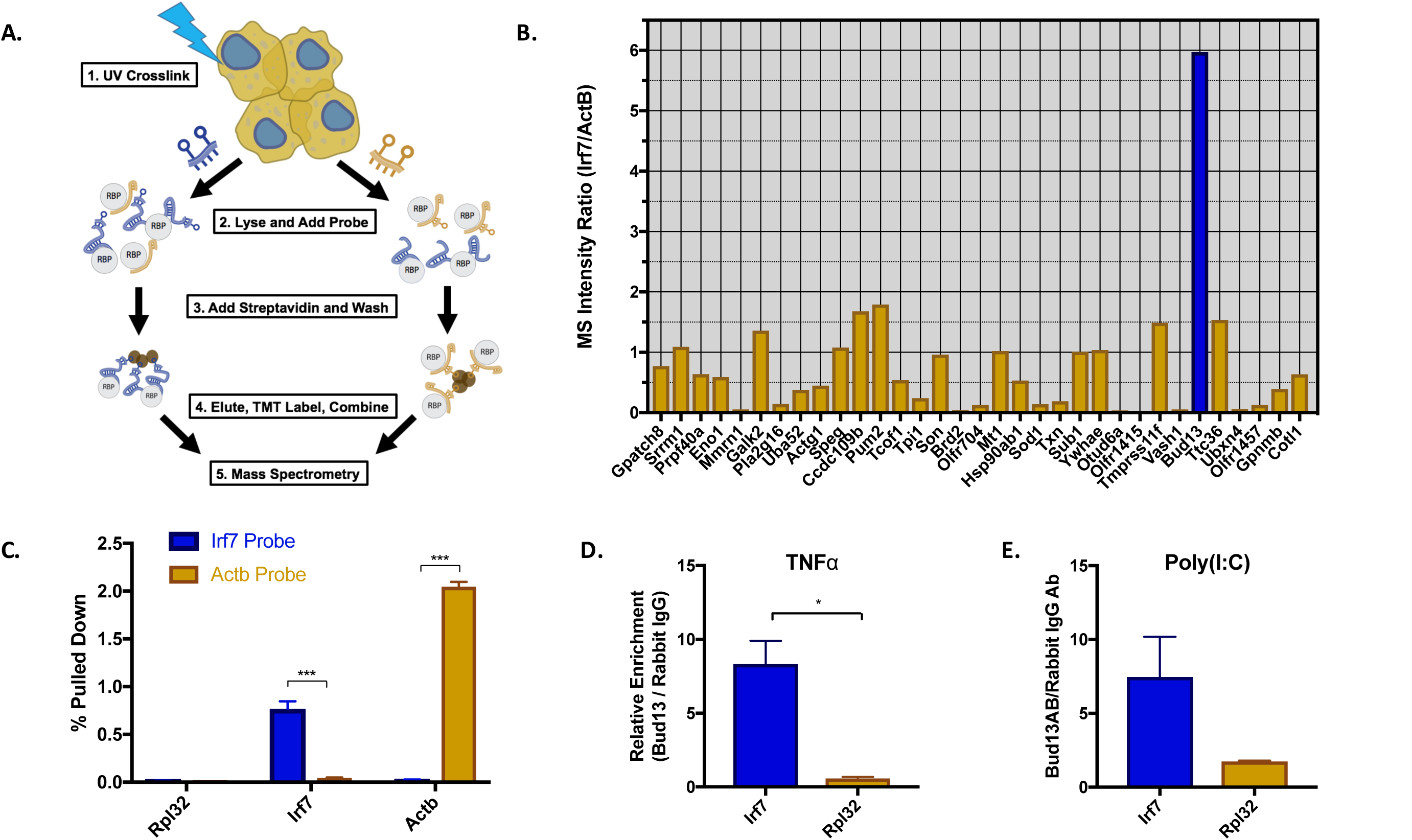
RAP-MS and RIP identifies Bud13 as an RNA binding protein that interacts with IRF7 mRNA. **(A)** Outline of the RAP-MS procedure used to identify RNA-binding proteins on transcritps of interest. **(B)** TMT ratio (Irf7/Actb) for proteins identified as enriched on either Irf7 (TMT ratio >1) or Actb (TMT ratio <1) transcripts. **(C)** qRT-PCR analysis of transcripts captured via RAP for Irf7 (blue) and Actb (gold) probes. **(D)** RIP followed by qRT-PCR for Irf7 and Rpl32 in TNFα stimulated BMDMs. Shown is the relative enrichment of transcripts captured in Bud13 RIP as compared to Rabbit IgG RIP. **(E)** Same as (d) except stimulation with poly(I:C). Data are representative of two independent experiments (**(C-E)**, mean + SEM). *P < 0.05, **P < 0.01 and ***P < 0.001 (t-test).

Following RAP-MS, we confirmed Bud13 enrichment on Irf7 transcripts by performing RNA Immunoprecipitation (RIP) followed by qPCR. Using formaldehyde cross-linked, BMDMs stimulated with TNFα for 30 minutes or Poly(I:C) for 12 hours, we observed >7-fold enrichment of *Irf7* transcripts associated with Bud13 immunoprecipitates as compared to Rabbit IgG control immunoprecipitates (Fig. 2D, E). In contrast, no differential enrichment of Rpl32 was observed. Thus, isolating the proteins associated with Irf7 mRNA transcripts led to the identification of Bud13, and immunoprecipitation of Bud13 protein confirmed enrichment of Irf7 mRNA.

### Bud13 knockdown leads to increased retention in the weak Irf7 intron

To determine whether the enrichment of Bud13 had an effect on Irf7 mRNA processing, we used an shRNA approach to knockdown Bud13 protein levels in BMDMs (Fig. S1A, B). To quantify differences in splicing between the shBud13 sample and the scrambled control sample, we calculated the difference in the previously mentioned splicing ratio (SR) metric between shBud13 and control for each junction at each time point. This resulting value was designated the ∆SR. A positive ∆SR indicates a junction is more unspliced in the shBud13 sample while a negative ∆SR indicates a junction is more unspliced in the control sample.

RNA-seq was performed on RNA from unstimulated BMDMs, as well as macrophages stimulated with TNFα for 0, 30, 60, and 120 minutes. Bud13 knockdown led to a further increased retention of the fourth intron in *Irf7* (Fig. 3A - highlighted intron). Further, the sequencing coverage plots showed little variation in splicing for the other seven introns in the transcript. This was confirmed when splicing was quantified using the ΔSR metric (Fig. 3B). At all stimulation time-points, the ∆SR value for the fourth intron was significantly greater than 0, indicating an increase in retention when Bud13 levels were reduced. In contrast, the ∆SR values for the other introns in the Irf7 transcript are negligible. It appears that Bud13 plays a specific role of aiding in the excision of the poorly spliced junction but is not required for total splicing of other introns in the transcript, at least as indicated by the partial knockdown with an shRNA. We next looked at how this retention affected the induction kinetics of Irf7. We observed that the increased retention seen upon knockdown of Bud13 led to decreased induction of Irf7 mRNA in response to TNFα stimulation (Fig. 3C, S2 C, D, E), consistent with the idea that intron retention leads to transcript degradation.^14^

**Figure 3:**
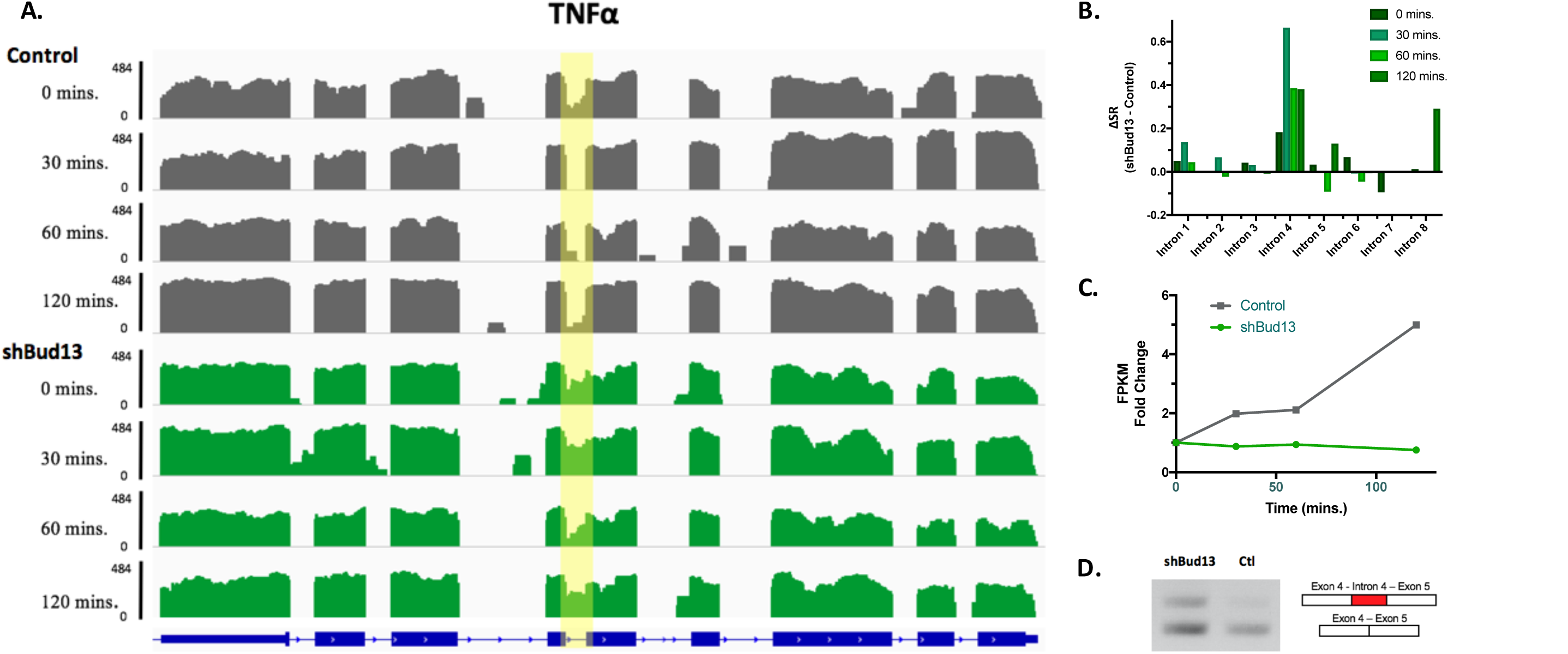
Bud13 knockdown leads to increased retention in the poorly splicing intron of Irf7. **(A)** Description of ∆SR metric used to calculate differences in splicing at individual time-points for given junctions. **(B)** Histogram of mapped reads corresponding to the TNFα-induced expression of Irf7. The poorly spliced fourth intron is highlighted. shBud1 3 samples are shown in green. Control samples are shown in grey. **(C)** ∆SRs calculated for each junction in the Irf7 transcript. Shade represents columns different stimulation time-point. **(D)** Irf7 FPKM fold change with respect to time stimulated. shBud13 is shown in green, control is shown in grey. **(E)** Splicing gel from RNA extracted from BMDMS stimulated for 30 mins. TNFα.

### Bud13 knockdown alters the type I interferon response

Because Irf7 is known as a ‘master regulator’ for robust type I interferon production^7^, we next investigated the effect of Bud13 knockdown on a type I interferon response. To do so, we stimulated BMDMs with the TLR3 agonist Poly(I:C) for up to 24 hours. Activation of TLR3 leads to the production of type I interferons followed by the downstream induction and activation of Irf7, which serves to amplify the type I interferon response via positive feedback^8^. We again observed differential splicing between the shBud13 time-points and the control time-points in intron 4 of Irf7 (Fig. 4A). Quantified using the ∆SR metric, five out of the six poly(I:C) time-points have high intron 4 ΔSR values (Fig. 4B). The ∆SR values for the other introns in the Irf7 transcript again fluctuate around 0, indicating that the fourth intron of Irf7 has a specific dependence on Bud13 for efficient splicing. As is the case with TNFα, this decrease in splicing efficiency alters Irf7 induction kinetics. Less Irf7 mRNA is induced at 240, 720, and 1440mins of poly(I:C) stimulation (Fig. 4C). This reduction in Irf7 mRNA leads to a decrease in nuclear Irf7 protein levels (Fig. 4D).

**Figure 4:**
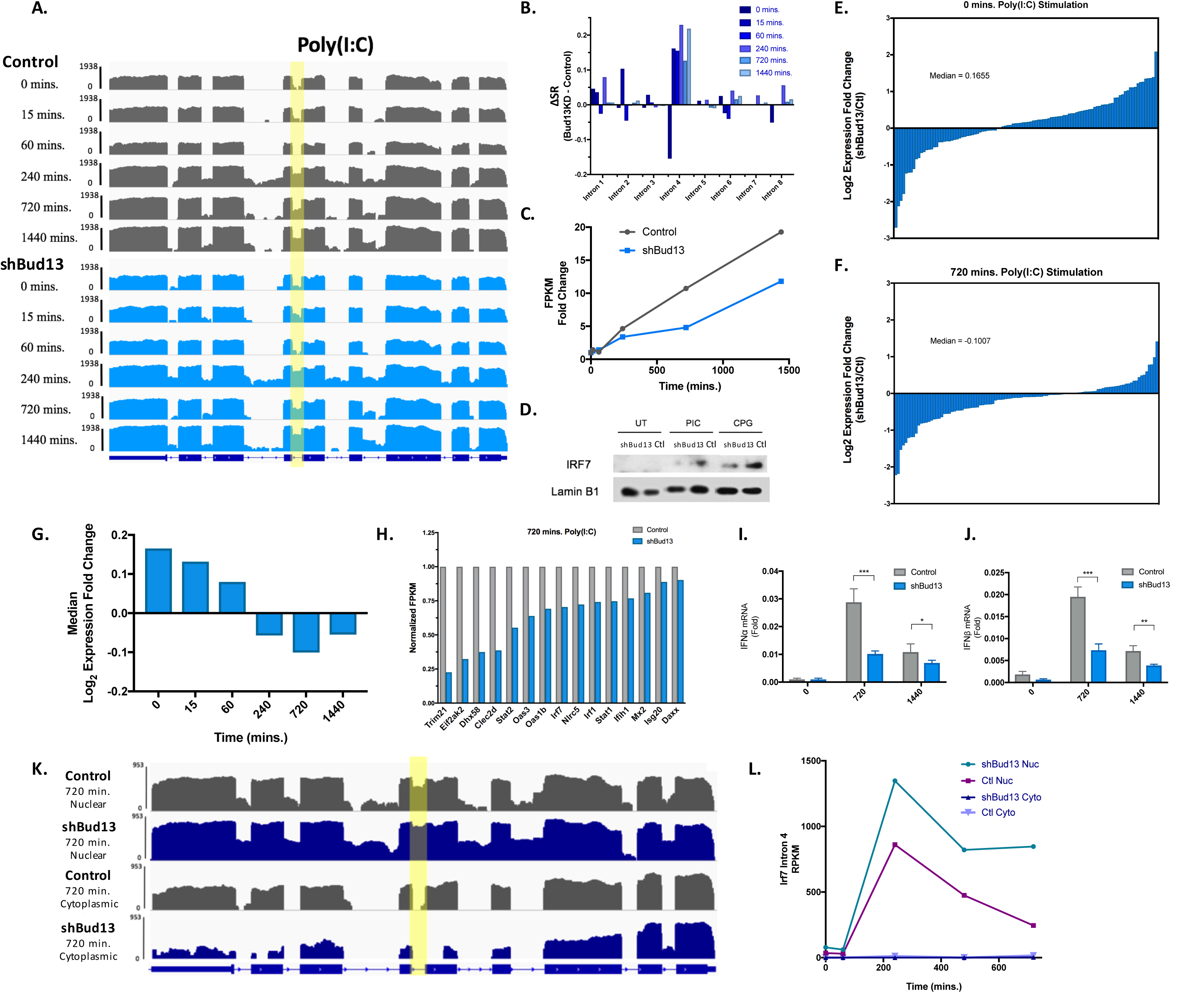
Bud13 knockdown alters the type I interferon response. **(A)** Histogram of mapped reads corresponding to the TNFα-induced expression of Irf7. The poorly spliced fourth intron is highlighted. shBud13 samples are shown in blue. Control samples are shown in grey. **(B)** ∆SRs calculated for each junction in the Irf7 transcript. Shade of columns represents different stimulation time-point. **(C)** Irf7 FPKM fold change with respect to time stimulated. **(D)** Immunoblot analysis of Irf7 protein after nuclear fractionation from BMDMs left untreated (UT) or treated with poly(I:C) (PIC) or CpG for 12h. Lamin B1 serves as loading control. **(E)** Log_2_ expression fold change (shBud13/control) for 119 ISGs in unstimulated BMDMs (median = 0.1655). **(F)** As in **(E)** for stimulated BMDMs (720 mins poly(I:C) (median = −0.1007). Wilcoxon rank-sum between **(E)** and **(F)**, *P*< .001. **(G)** Median log2 expression fold change (shBud1 3/control) for selected ISGs in unstimulated BMDMs, and BMDMs stimulated for 15, 60, 240, 720, and 1440 mins. (Wilcoxon rank-sum, *P<* .001, for any of the ‘early’ time-points (0, 15, 60 mins) compared to any of the ‘late’ time-points (240, 720, 1440 mins). **(H)** Normalized FPKM expression levels in shBud13 and control samples at 720 mins poly(I:C) stimulation for select ISGs. **(I)** RT-qPCR analysis of IFNα mRNA levels in unstimulated BMDMs and BMDMs stimulated with poly(I:C) for 720 mins and 1440 mins. **(J)** Same as **(I)** for IFNβ. **(K)** Nuclear fraction (top) and cytoplasmic fraction (bottom) histograms of mapped reads corresponding to the poly(I:C)-induced expression of Irf7 (720 mins). The poorly spliced fourth intron is highlighted. shBud13 samples are shown in blue. Control samples are shown in grey. Nuclear ∆SR = 0.35. **(L)** Nuclear and cytoplasmic FPKM for Irf7 intron 4 from fractionated BMDMs stimulated with poly(I:C). Data is representative of two independent experiments (i,j) and is represented as mean + SEM. * denotes p < 0.05, ** denotes p < 0.01, and *** denotes p < 0.001 using a Student’s t test. Results are presented relative to those of Rpl32 **(I,J)**.

Next we looked at how this reduction in Irf7 would alter the production of RNA from interferon signature genes (ISGs). Expression of 119 ISGs (selected based on upregulation in response to IFNα ^15^; see methods) was examined. In unstimulated BMDMs, used as a baseline, the median log_2_ expression fold change (FPKM shBud13/ FPKM control) is 0.1655 (Fig. 4E). In contrast, at 720 mins of stimulation, the median log_2_ expression fold change shifts to −0.1007 (Fig. 4F), indicating a significant decrease in ISG expression in the shBud13 sample compared to the control sample at this time-point compared to the baseline (Wilcoxon rank-sum, *P*< .001). This significant decrease in ISG expression remained true when comparing any of the ‘early’ time points (0, 15, 60 mins) to any of the ‘late’ timepoints (240, 720, 1440 mins) (Wilcoxon rank-sum, *P*<0.001). Dampened levels of Irf7 led to defects in the amplification of the type I interferon response, and a decrease in ISG expression later in the time-course (Fig. 4G). Trim21, Stat1, Stat2, Ifih1, and Dhx58 are among the notable ISGs with reduced expression at 720 mins in the shBud13 sample (Fig. 4H). Expression of both IFNα and IFNβ were significantly reduced at 720 and 1440 mins (Fig. 4I, J). To ensure differential expression of ISGs was not due to splicing defects from Bud13 knockdown, we quantified the ∆SR for every ISG junction at 720 mins. The fourth intron of Irf7 has the greatest ∆SR at 0.227. Only four other junctions of the 375 that were examined have ∆SRs greater than 0.1, and the majority of junctions have ∆SRs close to 0 (Fig. S3; mean = 0.002, median = 0). Similar results were obtained when BMDMs were stimulated with the TLR9 agonist CpG (Figure S4). Taken together, we conclude that Bud13 deficiency results in a highly compromised type I interferon response.

Finally, we examined whether Irf7 pre-mRNA with a retained fourth intron was able to exit the nucleus and enter the cytoplasm. BMDMs were stimulated with poly(I:C) and fractionated into a nuclear fraction (containing the nucleoplasm and chromatin) and a cytoplasmic fraction. RNA-seq was performed on RNA from the cytoplasmic fraction and the Irf7 mRNA was found to be almost completely spliced (Fig. 4K, L). In the nucleus, we again observed retention of Irf7 intron 4 (Fig. 4K, highlighted intron, ∆SR. = 0.35 at 720 mins). Taking the intronic reads across stimulation time-points, we notice both that the shBud13 nuclear sample is more unspliced and that the cytoplasmic samples in both the shBud13 and control samples are completely spliced. Thus, unspliced Irf7 is either being degraded in the nucleus, or it makes it to the cytoplasm and is degraded extremely quickly, such that virtually no signal can be detected via RNA-seq.

### Global analysis of the role of Bud13 in BMDMs

We next investigated global splicing differences caused by Bud13 knockdown. Using the TNFα stimulated data-set, ∆SR was calculated for every junction in every expressed gene. To determine whether splicing differences caused by Bud13 knockdown led to altered gene expression, we compared the effect of the Bud13 knockdown on genes that contained a Bud13 dependent junction to those that did not. (see methods). The median log_2_ expression fold change (FPKM shBud13/ FPKM control) for genes containing a Bud13 dependent junction was −0.5084. In contrast, the median log_2_ expression fold change (FPKM shBud13/ FPKM control) for genes without any junctions affected by Bud13 knockdown was −0.2170. Thus, we conclude there is an inverse relationship between IR due to Bud13 knockdown and gene expression (Wilcoxon rank-sum, *P*< .01) (Fig. 5A).

**Figure 5:**
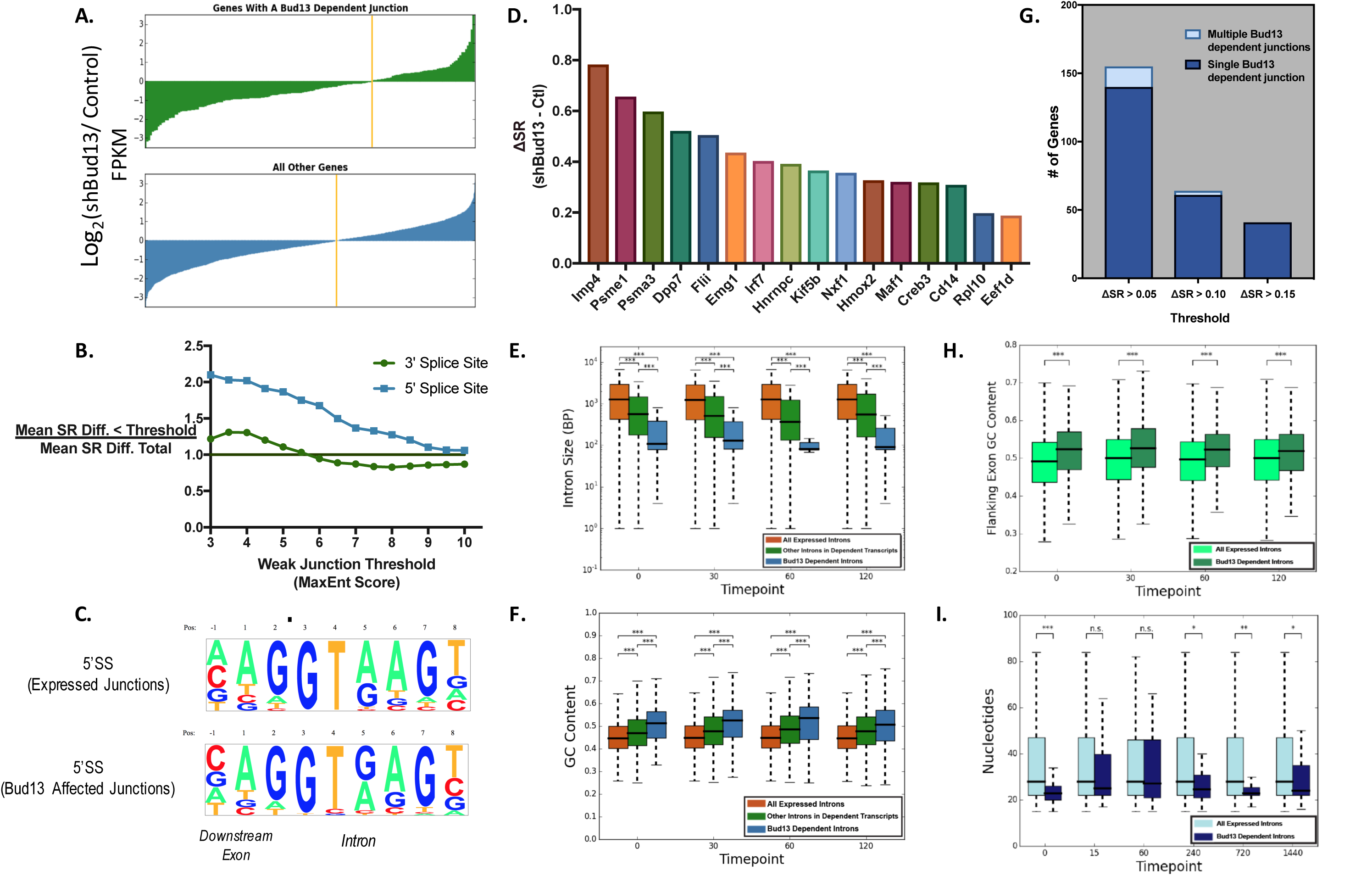
Global analysis of the role of Bud13. **(A)** Transcripts were classified as ‘Bud13 dependent’ if they had a junction with a ∆SR. >0.15. The log_2_ expression fold change (FPKM shBud13/ FPKM control) for each gene represented by the transcripts in the ‘Bud13 dependent’ category as well as all other genes is shown. Median ‘increased IR’ = −0.5084. Median ‘decreased IR’ = −0.2170. (Wilcoxon rank-sum, *P*< .01). **(B)** Mean ∆SR. for junctions below the indicated threshold (x-axis) vs. mean ∆SR. for all junctions. Threshold applied for the 5’ splice site (blue) and the 3’ splice site (green). **(C)** 5’SS motif for all expressed junctions as compared to junctions that show retention upon Bud13 knockdown (∆SR. > 0.15). (**D)** Ranked bar chart showing genes with a junction most affected by Bud13 knock-down in all samples during TNFα stimulation. See S7 for histograms relating to most affected junctions. **(E)** Size of intron for introns retained upon Bud13 knockdown (∆SR. > 0.15) (blue), in introns located in the same transcript as those affected by Bud13 (green), and in introns from all expressed transcripts (orange). **(F)** Same as **(E)** for GC content. (G) Grouped bar chart depicting the number of genes that have a single Bud13 affected junction vs. multiple Bud13 affected junctions using three different ∆SR thresholds. **(H)** Flanking exon GC content for exons that flank introns retained upon Bud13 knockdown (∆SR. > 0.15) (dark green) as compared to exons that flank introns from all expressed transcripts (light green). **(I)** Distance from the branch point to the 3’ splice site for introns retained upon Bud13 knockdown (∆SR. > 0.15) (dark blue) as compared to introns from all expressed transcripts (light blue). **(E, F, H, I)** data from BMDM TNFα stimulation. Box plots show median (center line), interquartile range (box) and tenth and ninetieth percentiles. \**P* < 0.05, \*\**P* < 0.01 and \*\*\**P* < 0.001 (Mann-Whitney *U*-test).

Next, it was of interest to us to identify sequence elements that led Bud13 to have its specific splicing effect. The most evident element to explore was the effect of splice site strength on Bud13-dependent splicing. Previous work has shown that the yeast orthologue of Bud13 plays a role in efficient splicing for a junction with a weak 5’ splice site^13^. Further, the junction affected in Irf7 has a non-consensus 5’ splice site. To investigate this issue, we first quantified every 5’ and 3’ splice site using a maximum entropy model.^12^ Then, we took progressively weaker splice site thresholds, and compared the mean ∆SR for every junction below that threshold to the mean ∆SR of every junction in the data-set (Fig. 5B). We saw that as the splice site threshold for the 5’ splice site became progressively weaker, the mean ∆SR for junctions weaker than that threshold increased and thus there was a greater Bud13 splicing effect. This result was not seen when the same analysis was applied to the 3’ splice site. We then analyzed the Bud13 splicing effect with respect to other features known to correlate with IR.^6^ Across all time-points for both TNFα (Fig. 5E, F, H, I) and Poly(I:C) (Fig. S7 A-C), Bud13 dependent introns were dramatically smaller and had increased G/C content in both the intron and in the flanking exons. We also noticed that the distance from the branch point to the 3’ splice site was smaller in the Bud13 dependent introns than in the total data-set (Fig. 5I, Fig. S7 D). This could be a byproduct of the smaller intron length; however, it is of interest because Bud13 has been shown in yeast to bind just downstream of the branch point.^16^ A significant difference was not seen in branch point strength and Bud13 splicing effect (Fig. S7 E, F). Of note, the fourth intron of Irf7 is among the most Bud13 dependent junctions in both the TNFα and Poly(I:C) data-sets (Fig. 5D., Fig. S7H, see methods for analysis details). Further, almost all transcripts contain only a single Bud13 dependent junction, even when low thresholds are used to quantify dependency (Fig. 5G.). This is again similar to what we observed in Irf7.

Finally, using enhanced crosslinking and immunopreciptation (eCLIP)-seq data from the ENCODE Project Consortium,^17^ we investigated Bud13 binding specificity across the genome. We found that in K562 and Hep G2 cells, the majority of Bud13 eCLIP-sequencing reads were located downstream of the branchpoint near the 3’ splice site (Fig. 6A, B), consistent with what is seen in yeast.^16^ There is some read density near the 5’ splice site, which we hypothesize is due to Bud13’s association with the spliceosome. Although it may not bind near the 5’ splice site, factors in the spliceosome that interact with Bud13 may immunoprecipitate with Bud13, leading to a 5’ signal. Data for Sf3b4 and Prpf8, known RBPs that interact with the 3’ and 5’ splice site respectively, is also shown (Fig. 6A, B). After peak calling, we counted the location of significant peaks. The majority of peaks are in intronic regions or intron-exon junctions. Further, of the peaks that lie in intron-exon junction, most are located at the 3’ junction (Fig. 6C, D). Lastly, when comparing introns that have an overlapping eCLIP peak to all introns from expressed transcripts, we see both a length and G/C% bias (Fig. 6E, F). Bud13 peaks tend to fall in smaller introns that are GC rich, a finding consistent with the ∆SR data.

**Figure 6:**
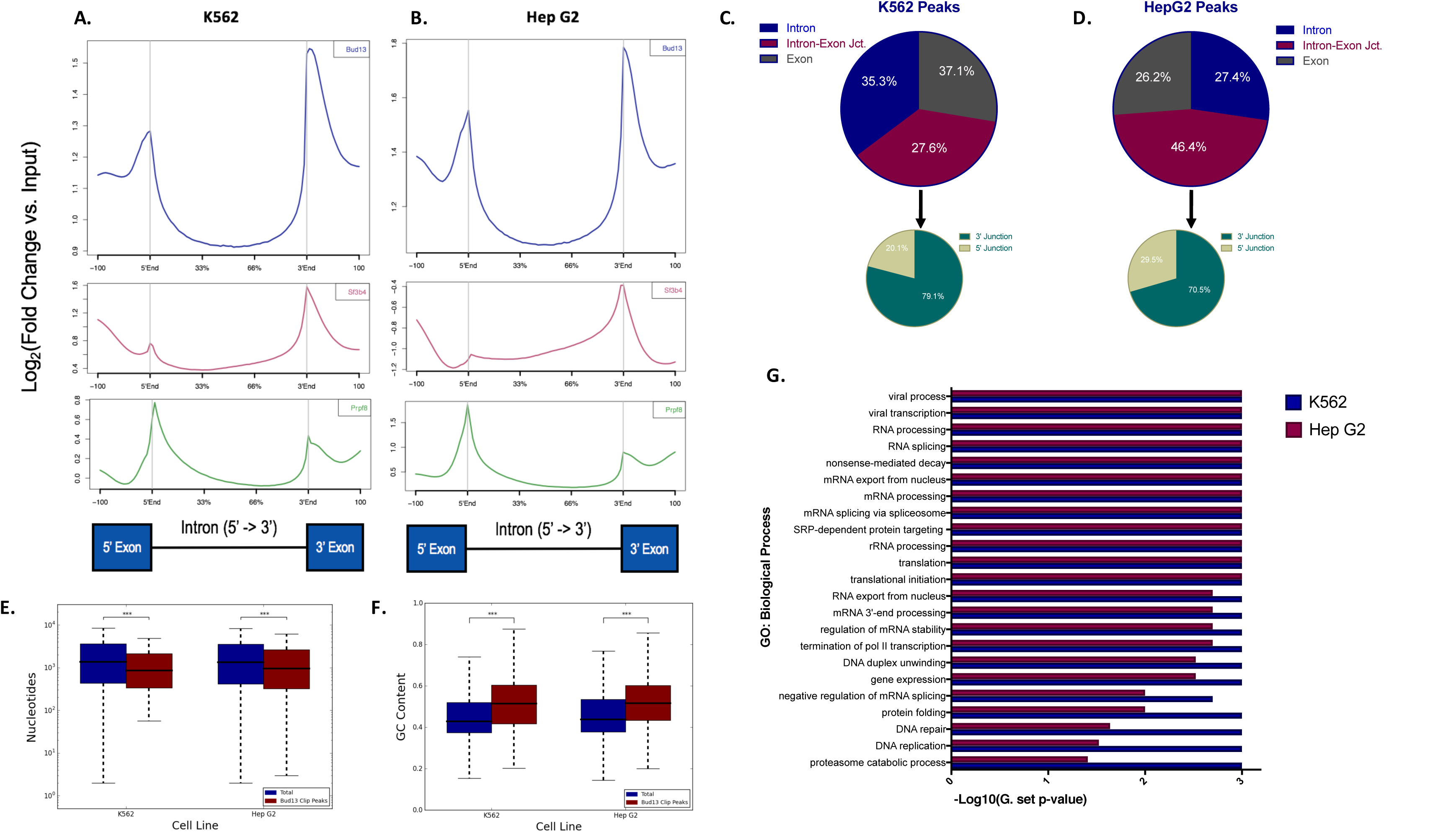
Bud13 interacts primarily near the 3’ splice site of small, GC rich introns. **(A)** eCLIP-seq read density plots in K562 cells. Bud13 eCLIP-seq is shown in blue (top), Sf3b4 is shown in maroon (middle), and Prpf8 is shown in green (bottom). **(B)** Same as in **(A)** but for Hep G2 cells. **(C)** Bud13 eCLIP-seq peak distribution. Peaks fell within either intronic regions, intron-exon junctions, or exonic regions. Peaks that fell within intron-exon junction were further classified as 5’ junction peaks or 3’ junction peaks (bottom). **(D)** Same as **(C)** but for Hep G2. **(E)** Size of all introns in expressed transcripts for the given cell line (dark blue) vs size of introns with overlapping eCLIP peak (maroon). Shown in K562 (left) and Hep G2 (right) cells. Box plots show median (center line), interquartile range (box) and tenth and ninetieth percentiles (whiskers). \**P* < 0.05, \*\**P* < 0.01 and \*\*\**P* < 0.001 (Mann-Whitney *U*-test). **(F)** Same as **(E)** for GC content. **(G)** GO terms (biological process) enriched among Bud13 eCLIP peaks in K562(dark blue) and Hep G2 (maroon) cells.

### Bud13 knockdown alters the BMDM response to VSV

Vesicular stomatitis virus (VSV) is a (-)ssRNA virus known to induce type I IFN through TLR7^18^. To test whether impairment of Irf7 due to Bud13 knockdown was present in VSV stimulated BMDMs, we infected both shBud13 and control BMDMs at an MOI of 5 and 10. At all time-points throughout infection in both MOIs, there was dampened Irf7 induction (Fig. 7A, B) as quantified by Taqman qPCR. Next, in order to determine the consequences of impaired Irf7 induction, we determined the yield of virus from BMDMs following a period of infection with a given input MOI. shBud13 BMDMs produce >10 times the amount of VSV as compared to control BMDMs (Fig. 7C). This difference in viral production is presumably due to the dampened type I interferon response, as well as the corresponding deficiency in ISG production associated with Bud13 knockdown. Finally, we notice increased infectivity across a 24-hour time-course, a finding again consistent with our previous data showing that Bud13 alters Irf7 splicing and thus hinders its ability to amplify the type I interferon response (Fig. S8).

**Figure 7:**
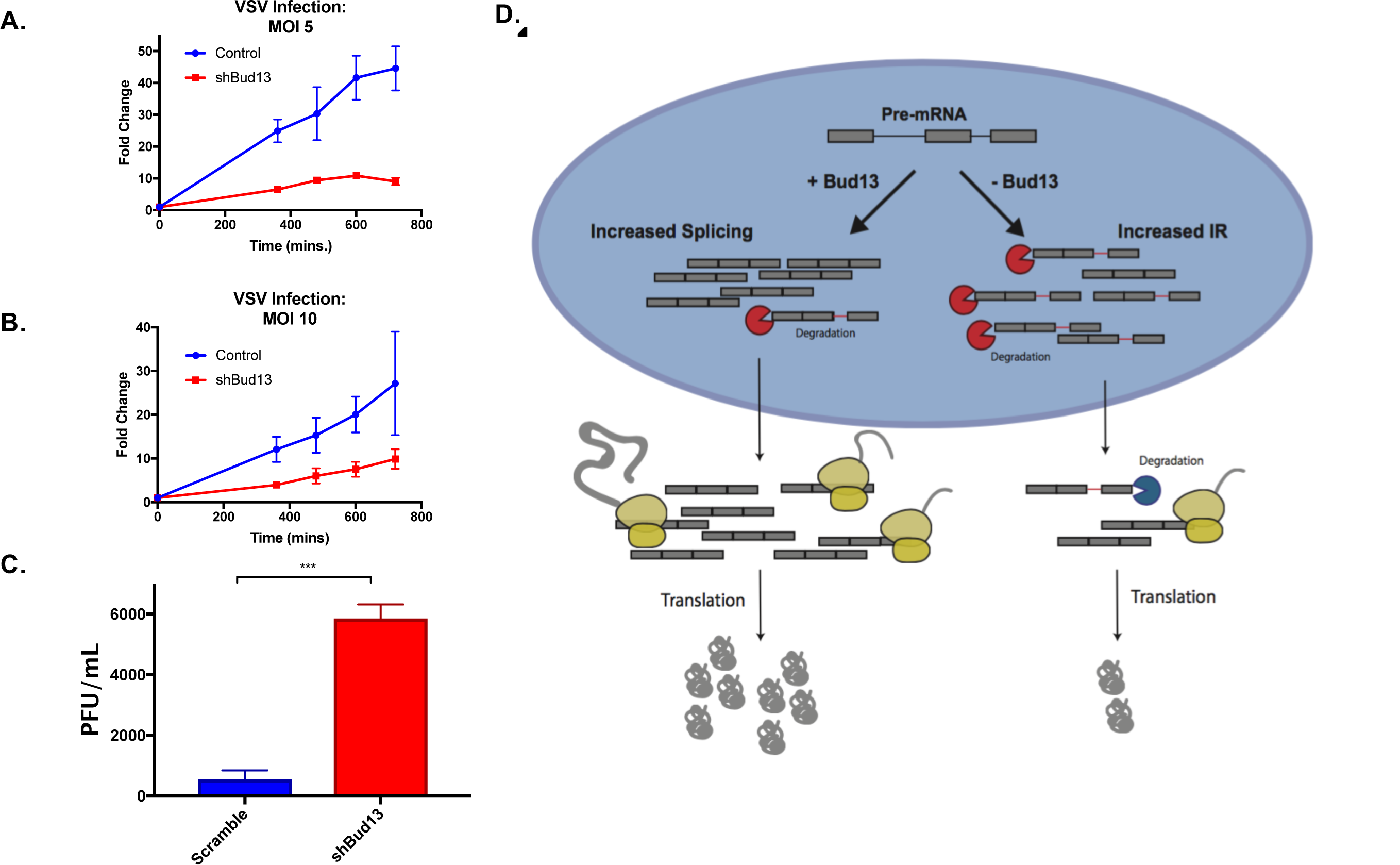
Bud13 knockdown alters the BMDM response to VSV. **(A)** RT-qPCR analysis of Irf7 mRNA levels in infected control or shBud13 BMDMs stimulated with VSV (MOI 5) across 24 hours. **(B)** Same as in **(A)** except stimulated at an MOI of 10. Results are presented relative to those of Rpl32. **(C)** PFU/mL for viral supernatant from infected control or shBud13 BMDMs. Data is representative of two **(A, B)** or three independent experiments **(C)** and is shown as mean + SEM. * denotes p < 0.05, ** denotes p < 0.01, and *** denotes p < 0.001 using a Student’s t test

## DISCUSSION

In this study, we sought proteins that might relate to the poor splicing of an intron in Irf7 transcripts. Using RAP-MS, we identified Bud13 as a protein that has the ability to increase splicing of the Irf7 intron. In the absence of *Bud13*, in response to inflammatory stimulus, macrophages produced Irf7 with increased intron retention (IR) and notably less mature Irf7 transcript and protein (Fig. 3C, 4C, 4D, S4C). Irf7 is the interferon-inducible master regulator of the type-I interferon-dependent immune response.^7^ Correspondingly, depletion of Bud13 led to a general reduction in ISG and cytokine production, implying a compromised type I interferon response (Fig. 4E-J, S4D-G). This splicing and corresponding expression defect upon Bud13 depletion was observed under various stimulation regimens and times. We found that macrophages deficient for Bud13 were strikingly more susceptible to infection by VSV, presumably owing to the reduction in Irf7 transcript levels (Fig. 7).

We observed the Bud13 splicing dependence in other introns of other genes. A number of short, GC-rich introns with non-consensus splice donor sites were excised inefficiently when Bud13 levels were depleted (Fig. 5). As was the case with Irf7, this increased IR reduced mature transcript levels (Fig. 5A). Transcripts containing retained introns have been shown in the literature to be degraded by two mechanisms: (1) nuclear degradation via the RNA exosome, (2) cytoplasmic degradation upon detection of a pre-termination codon (PTC) via the NMD decay machinery. Although the majority of these introns contain a PTC, it remains to be determined whether degradation is occurring in the nucleus or cytoplasm.^14^ ^19^

Bud13 was originally identified as a part of a “Retention and Splicing” (RES) complex^13^ in yeast. However, yeast Bud13 (ScBud13) and mammalian Bud13 are significantly different lengths (266 vs. 637 amino acids) ^20^, with only the mammalian protein containing a large, disordered arginine-rich N-terminal domain. ScBud13 counteracts IR in introns within the mediator complex, mating genes, and tRNA modifying genes^21-23^, which in turn impair yeast budding. In connection with the RES complex, ScBud13 is thought to safeguard formation of the ‘B^act^ complex’ of the spliceosome.^24^ In the B^act^ stage, the 5’ splice donor and branch point are recognized by the spliceosome. However, progression to catalysis of the first step of the splicing reaction requires remodeling of several spliceosome components.^25^ Lack of the RES complex has been shown to lead to premature binding of Prp2, a quality control factor that is responsible for spliceosome remodeling as well as the disassembly of suboptimal substrates. It has been hypothesized that ScBud13 and the RES complex temporally regulate Prp2 binding.^24^ In the mammalian context, short, GC-rich introns with weak donor sites may be particularly susceptible to Prp2-mediated disassembly, which may explain the specificity of IR events upon Bud13 depletion.

In yeast, differential studies using mass spectrometry^26^ and cross-linking have established that some ScBud13 is detectable in preparations of stalled B, B^act^, and B* complexes. One cryo-EM structure of the yeast spliceosome found density corresponding to ScBud13 in a stalled B^act^ pre-catalytic complex, although a structure of the stalled B complex found only weak density for ScBud13.^22,27^ In mammals, structural evidence of Bud13 is limited. Given the partial sequence homology between all members of the yeast RES complex and their mammalian counterparts, it is not surprising that Bud13 (and other RES complex members) are often undetectable in preparations of stalled spliceosomes using cross-linking and mass-spectrometry. Furthermore, Bud13 was not detected in a recent human cryo-EM structure of a stalled B complex.^28^ Taken together, it is not yet possible to determine if the sub-stoichiometric nature of Bud13 in mammalian spliceosome complexes is because it is constitutively associated but highly transient or because it serves as a non-essential accessory to spliceosome function. Cryo-EM studies, as well as single molecule studies, would seem to suggest compositional heterogeneity of the spliceosome, and that the Bud13-endowed spliceosome may catalyze the splicing reaction in a fundamentally different way than is used in its absence.^24,29,30^

Recently, Bud13 in zebrafish was shown to regulate levels of IR in short, GC-rich introns in knockout studies.^31^ Indeed, both in Zebrafish^31^ and C. Elegans,^32^ Bud13 deficiency has been reported to lead to embryonic lethality. Our results show that mammalian Bud13 shares this splicing fidelity function, and deficiency may prevent proper development. Despite this, knockdown and knock-out cell lines have displayed no overt growth defects, suggesting a developmental but not immune-cell intrinsic dependence on Bud13 for survival.

Bud13 might simply represent a mechanism that evolved to counter intron retention in a subset of introns that require splicing but happen to be inherently weak. Furthermore, as previously mentioned, IR has emerged as a post-transcriptional mechanism used by the cell to fine-tune gene expression. It is possible that the presence of these weak introns, coupled with regulation of Bud13, aid in the fine-tuning of the transcriptome. Mammalian Bud13 undergoes intensive phosphorylation and acetylation.^20^ Additionally, it is intrinsically disordered and co-localizes with nuclear speckles.^33^ Alterations in the levels of Bud13 or regulation of post-translation modifications that could affect Bud13 activity or lead to speckle sequestration might allow a cell the ability to quickly adjust the expression of a subset of transcripts.

In summary, we found that Bud13 modulates gene expression through its ability to alter IR, often in notably small, GC-rich introns with weak splice sites. Deficiency of Bud13 results in IR and concomitant decreased gene expression in transcripts such as Irf7, dampening the type I interferon response and increasing viral susceptibility. Therefore Bud13, in mediating Irf7 gene expression, presents a potential therapeutic target for the treatment of infections or autoimmune conditions. Future studies should seek to understand why Bud13 is vital for the efficient splicing of only a subset of junctions and whether or not this junction specificity plays an active role in regulating gene expression. If modulated, this strategy by which components associated with the spliceosome rescue transcripts from intron retention and degradation may represent a previously underappreciated layer of regulation in many gene expression programs.

## ACKNOWLEDGEMENTS

The authors would like to thank Mario Blanco and Mitchell Guttman (Dept. of Biology, Caltech) for assistance with eCLIP analysis; and Logan Vlach, Patricia Turpin, Mati Mann and Alok Joglekar (Dept. of Biology, California Institute of Technology) for experimental and computational assistance. Additionally, the authors would like to thank Genhong Cheng (Department of Microbiology, Immunology & Molecular Genetics, UCLA) for VSV and Jae Jung (Department of Molecular Microbiology & Immunology, USC) for VSV-GFP. This work was funded from a grant from NIH (5R21AI126344) and from an endowment provided by the Raymond and Beverly Sackler Foundation.

## AUTHOR CONTRIBUTIONS

L.S.F, D.M., and D.B., conceived and designed experiments. L.S.F. conducted experiments. C.B. helped develop RAP-MS and knockdown experiments. L.S.F. and D.M. analyzed sequencing data. A.M. oversaw mass spectrometry and M.J.S. performed mass spectrometry analysis. L.S.F, D.M., and D.B wrote the manuscript with input from all authors.

## STAR METHODS

### Contact for Reagent and Resource Sharing

Further information and requests for resources and reagents should be directed to and will be fulfilled by the Lead Contact, David Baltimore.

### Experimental Model and Subject Detail

#### Animals

The California Institute of Technology Institutional Animal Care and Use Committee approved all experiments. C57BL/6 WT mice were bred and housed in the Caltech Office of Laboratory Animal Resources (OLAR) facility. C56BL6/J mice were sacrificed via CO2 euthanasia and sterilized with 70% ethanol. Femur and tibia bones harvested and stripped of muscle tissue. Bone marrow cells were resuspended in 20mL of fresh DMEM. 2.5 *10^6^ bone-marrow cells plated in a 150mm dish in 20mL of BMDM Media (DMEM, 20% FBS, 30% L929 condition media, and 1% Pen/Strep) and grown at 5% CO_2_ and 37°C. BMDM media was completely replaced on day 3 as well as a supplemental addition of 5mL L929 condition media on day 5.

### Cell Culture

Human embryonic kindey cells (HEK293T) from ATCC were cultured in DMEM supplemented with 10% FBS and 1% Pen/Strep. Cell line was maintened at 37°C in 5% CO_2_.

### Method Detail

#### Knockdown Experiments

BMDMs for knockdown experiments were grown as described above with a few additions. On days 3 and 4, retrovirus encoding shRNAs were added to cells. On day 5, cells were selected with puromycin (5ug/mL). On day 8, following ~72 hours of puromycin treatment, media was removed and 10mL of PBS w/ 2mM EDTA was added. Cells were lightly scraped and replated in either 6 well plates or 10-cm dishes depending on the experiment. Cells were left in BMDM media overnight. The following day, cells were stimulated with either 20ng/mL of TNFα, 5ug/mL Poly(I:C) (Sigma), or ODN 1585 (1-5 μM) (InvivoGen).

#### RNA Isolation

Total RNA was purified from BMDMs using TRIzol reagent (Ambion) as per the manufacturer’s instructions. Genomic DNA in RNA purifications was eliminated through treatment with Turbo DNase (Thermo Fisher Scientific) for 30 min at 37°C. 0.1-1μg RNA and 1μM dT(30) oligo (d14-954: 5’-AAGCAGTGGTATCAACGCAGAGTACT(30)) was heated at 80°C for 2.5min followed by snap cooling on ice. 10μL template-switch RT mix added (10μM template-switch oligo (TSO: 5’-AAGCAGTGGTATCAACGCAGAGTACACArGrGrG), 20mM DTT, 2X ProtoScriipt II Reverse Transcriptase Reaction Buffer (NEB), 1mM dNTPs, 40U Murine RNAse Inhibitor (NEB), and 200U ProtoScript II (NEB) Reverse Transcriptase. Reaction incubated in thermocycler with the following program: 1. 42°C for 30min, 2. 45°C for 30min, 3. 50°C for 10min, followed by deactivation of RT for 10min at 80°C.

#### RNA Fractionation

Confluent 10-cm dish of mature BMDMs were scraped into 400μL cold NP-40 lysis buffer, APJ1 (10mM Tris-HCl (pH 7.5), 0.08% NP-40, 150mM NaCl). Lysed cells layered onto 1mL cold sucrose 322 cushion, APJ2 (10mM Tris-HCl (pH 7.5), 150mM NaCl, 24% w/v sucrose) and centrifuged for 10min at 4°C and 13000 rpm. The supernatant from this spin represents the cytoplasmic RNA fraction, which is immediately added to 3 volumes of 100% ethanol and 2 volumes of buffer RLT (4M GuSCN, 325 0.1M β-mercaptoethanol, 0.5% N-lauroyl sarcosine, 25mM Na-citrate, pH7.2) and stored at −80°C until ready to purify RNA. Pellet, containing intact nuclei, is resuspended in 500μL TRIzol reagent. If the pellet was difficult to dissolve, it was heated at 50°C with occasional vortexing. 100μL chloroform added and shaken vigorously for 15-20s; allowed to phase separate at room temperature for 5min. Tube centrifuged at 4°C and 12000 x g for 15min. Clear upper aqueous phase removed to a new tube, ensuring white DNA mid-phase is not removed, and is immediately added to 3 volumes of 100% ethanol and 2 volumes of buffer RLT and stored at −80°C until ready to purify RNA. RNA is purified according to Qiagen RNeasy column protocol and eluted in 30μL nuclease-free H2O. RNA samples are DNAse treated with Turbo-DNAse and stored at −80°C.

#### Library preparation and RNA-Seq Analysis

Limited PCR amplifications was performed prior to library preparation. PCR reaction done with KAPA HiFi HotStart 2x ReadyMix, 5% cDNA, and 1μM primer (d14-955: 5’-AAGCAGTGGTATCAACGCAGAGTACT). Thermal cycler programmed for 120 seconds at 95°C as initial denaturation, followed by 14 cycles of 30sec at 95°C for denaturation, 30sec at 62.5°C as annealing, 150sec at 72°C for extension, and final extension at 72°C for 5 min. PCR reactions 0.9X SeraMag and eluted in 25μL Concentrations of purified library determined using Qubit High Sensitivity dsDNA kit (Invitrogen) as described. Full length cDNA libraries were barcoded using the Nextera XT Tagmentation protocol (Illumina).

#### RNA-Antisense Purification

RNA antisense purification-mass spectrometry (RAP-MS) was performed as described in McHugh et al. with a few alterations. Briefly, we designed three 90-mer DNA oligonucleotide probes that were antisense to the complementary target RNA sequence in both *Irf7* and *Actb* transcripts. Each probe was targeted to a different location on the transcript and modified with a biotin in order to enable capture of DNA:RNA hybrids on streptavidin coated magnetic beads.

*RNA Prep and Lysis:* ~250million cells, or 25 150mm plates of BMDMs were used for each capture. Following stimulation with TNFα (20ng/ml) for 30 minutes, ~5-10 mL of PBS w/ 2mM EDTA was added to each plate and cells were removed by lightly scraping. Cells were pelleted, resuspended in PBS, and poured into a new 150mm plate. The cells were then crosslinked in Spectrolinker at 254 nm wavelength with 0.8 J/cm2 (instrument setting: 8000 x 100 uJ/cm^2^). Following crosslinking, cells were again pelleted, at which point the pellet could be frozen and stored at −80°C. Cells were lysed in 2mL of lysis buffer per capture (10 mM Tris pH 7.5, 500 mM LiCl, 0.5% Triton X-100, 0.2% sodium dodecyl sulphate, 0.1% sodium deoxycholate) supplemented with Protease Inhibitor Cocktail (EMD Millipore) and 1000 U of Murine RNase Inhibitor (New England Biolabs). We found the smaller the volume used per sample, the more efficient the capture was downstream and thus the minimum volume needed to lyse cells should be optimized. Samples were incubated for 10 min on ice to allow lysis. Following lysis, sample was passed through 20-gauge needle once and then 26-gauge needle 3-5 times to disrupt the pellet and shear genomic DNA. In between passing the sample through the 26-gauge needle, the sample was sonicated on ice with a microtip set at 5W power for a total of 30 s in intermittent pulses (0.7 s on, 1.3 s off). Samples were then mixed with twice the lysate volume of 1.5x LiCl/Urea Buffer (the final buffer contains 10 mM Tris pH 7.5, 500 mM LiCl, 0.5% Triton X-100, 0.2% SDS, 0.1% deoxycholate, 4 M urea). Lysates were incubated on ice for 10 min then cleared by centrifugation for 10 min at 4,000g.

*Pre-clearing lysate:* BioMag streptavidin beads (Bang Laboratories Inc.) were first washed 3x in 0.25-0.5ml of 500mM LiCl/4M Urea buffer (10 mM Tris pH 7.5, 500 mM LiCl, 0.5% Triton X-100, 0.2% SDS, 0.1% deoxycholate, 4 M urea). 50ul of beads were added to each sample and the samples were incubated at 37°C for 30 min with shaking. Streptavidin beads were then magnetically separated from lysate samples using a magnet. The beads used for preclearing lysate were discarded and the lysate sample was transferred to fresh tubes twice to remove all traces of magnetic beads. Input for quality control experiments can be removed at this point.

*Hybridization, Capture of Probes and Elution of Associated Protein:* Following pre-clearing, the biotinylated 90-mer DNA oligonucleotide probes specific for the RNA target of interest (vary per sample but ~5ul of 25uM per probe) were heat-denatured at 85°C for 3 min and then snap-cooled on ice. Probes and pre-cleared lysate were mixed and incubated at 55°C with shaking for 2 h to hybridize probes to the capture target RNA. 500mL of washed streptavidin beads (Bang Laboratories Inc.) were then added to each sample at 55°C with shaking for 30 mins. Beads with captured hybrids were washed 6 times with LiCl/Urea Hybridization Buffer. If needed, 1% of the beads can be removed for qPCR quality control experiment. TRIzol reagent can be added directly to beads to elute RNA. Beads were then resuspended in Benzonase Elution Buffer (20 mM Tris pH 8.0, 2 mM MgCl2, 0.05% NLS, 0.5 mM TCEP) and 125 U of Benzonase nonspecific RNA/DNA nuclease was added. Incubation occurred for 1-2 h at 37°C. Beads were then separated from the sample using a magnet. Supernatant was collected. Contaminant beads were removed by 5 rounds of magnetic separation on supernatant. Protein was precipitated overnight at 4°C with 10% trichloroacetic acid (TCA). TCA treated protein elution samples were pelleted by centrifugation for 30 min at 20,000g, then washed with 1 ml cold acetone and recentrifuged. Final protein elution pellets were air dried to remove acetone, resuspended in fresh 8 M urea dissolved in 40 ml of 100 mM Tris-HCl pH 8.5, and stored at −20°C.

*Mass Spec Prep. and Analysis* Performed as in McHugh et al. with few exceptions. Instead of SILAC we label proteins at the mass spec prep step using TMT (Thermo). After desalting on a Microm Bioresources C8 peptide MicroTrap column and lyophilization of peptide fraction, lyophilized protein pellets were resuspended in 100mM TEAB at a concentration of 1ug/ul. We then added 1.64ul of TMT labelling reagent to each ug of sample. The reaction was incubated for one hour at room temperature. The reaction was quenched with 0.32ul of 5% hydroxylamine per ug of protein used and incubated for 15 mins at room temperature. Following quenching, the samples were mixed, desalted as before, lyophilized, and mass spec was performed on Orbitrap Fusion mass spectrometer using a TMT instrument method as described in Liu et al.^38^

Raw files were searched using MaxQuant (v. 1.5.3.30) against the UniProt mouse database (59550 sequences) and a contaminant database (248 sequences). TMT 6plex was selected as the quantitation method with a reporter mass tolerance of 0.3. Oxidation of methionine and protein N-terminal acetylation were variable modifications and carbamidomethylation of cysteine was fixed modification. A 1% protein and peptide false discovery rate as estimated by the target-decoy approach was used for identification.

#### RNA Immunoprecipitation

RNA immunoprecipitations were performed as previously described. Between 5-10 confluent 15 cm^2^ dishes of BMDMs per sample were stimulated with either 20ng/mL of TNFα for 30 minutes or 5ug/mL Poly(I:C) for 12 hours. Following stimulation, proteins were cross-linked to DNA by adding formaldehyde directly to the media to a final concentration of 0.75%, with light shaking at room temperature for 10 mins. To quench the crosslinking reaction, glycine to a final concentration of 125 mM was added to the media and incubated with shaking for 5 mins at room temp. Media was then aspirated and cells were rinsed twice with 10 mL of cold PBS. Following the second wash, cells were scraped into 10mL of PBS and spun down gently (5 min, 4°C, 1,000xg). Final cell pellet was resuspended in 0.1-1mL of polysome lysis buffer (100 mM KCl, 5 mM MgCl_2_, 10 mM HEPES (pH 7.0), 0.5% NP40, 1 mM DTT, 100 U.ml RNase Inhibitor (NEB)) supplemented with Protease Inhibitor Cocktail (EMD Millipore). At this point the mRNP lysate was frozen. If needed, passing the lysate through a small gauge needle can help with lysate. Protein-G beads were pre-treated at 4°C with NT2 (50 mM Tris-HCl (pH 7.4), 150 mM NaCl, 1 mM MgCl_2_, 0.05% NP40) supplemented with 5% BSA to a final ratio of 1:5 for at least 1h before use. Appropriate amount of antibody per sample (optimized based on antibodyused but typically ~1-1 0ug) was added ot 250-500ul of bead/BSA slurry and incubated at 4°C. Following incubation, beads were spun down and washed with 1 ml of ice-cold NT2 buffer 4-5 times. Following final wash, beads were resuspended in 850ul of NT2 and supplemented with 200U of RNase inhibitor, 10 μl of 100 mM DTT and EDTA to 20 mM. Frozen lysate was thawed and centrifuged at 15,000*g for 15 mins. The cleared supernatant was removed and 100ul was added to the prepared beads. Input removed at this step. Beads and lysate were incubated for 4h at 4°C with mixing. The beads were washed 4-5 times with ice-cold NT2 and then resuspended in 100ul of NT2 buffer. 4ul of 5M NaCl was added incubated with shaking at 65°C for 2 hours. NT2 buffer can also be supplemented with 30 μg of proteinase K to release the RNP component. RNA was isolated by adding TRIzol reagent (Ambion) as per the manufacturer’s instructions. RNA was reverse transcribed and quantification was performed using TaqMan qPCR.

#### Immunoblot

BMDM samples were prepared as described previously. On day 8, they were plated at similar density. Following adherence, BMDMs were stimulated with Poly(I:C) or CpG for the indicated time. Cells were scraped into subcellular fractionation buffer (20mM HEPES (pH 7.4), 10 mM KCl, 2 mM MgCl_2_, 1mM EDTA, 1 mM EGTA). The cells were then passed through a 27 gauge needle 10 times, incubated on ice for 10 mins, and spun down at 720xg for 5 min. The pellet contained the nuclei, which was washed with fractionation buffer, passed through a 25 gauge needle 10 times, and centrifuged again at 720xg for 10 mins. The resulting pellet was resuspended in RIPA lysis buffer. Queal amounts of proteins were analyzed by immunoblot using the following reagents: anti-IRF7 (Millipore, ABF130), anti-Lamin B1 HRP conjugate (Cell Signalling, D9V6H), and anti-rabbit IgG HRP conjugate (Cell Signalling).

#### Viral Plaque Assays

Plaque assays were done one Vero cells. 2.5*10^5^ vero cells were plated in a 12 well plate the night before infection. Prior to infection, cells were checked to ensure confluence. VSV was serially diluted and infected in 12 well plate for 1 h. VSV was then removed and cells were layered carefully with DMEM supplemented with 2% FBS and 0.4% agarose. Plate was incubated for 2 days, and then fixed with 10% formaldehyde, for 1 h to overnight. Finally, agarose plugs were removed carefully and cells were stained with crystal violet.

#### VSV-GFP Infection Experiment

BMDMs were grown as described above in 150mm dishes. On day 8, following ~72 hours of puromycin treatment, media was removed and 10mL of PBS w/ 2mM EDTA was added. Cells were lightly scraped and 250,000 cells/well were replated in 12 well plates in BMDM media. Cells were left for 12 hours to adhere. Following adherence, VSV-GFP was added at the specified MOI for the specified amount of time. Following the time-course, cells were lightly scraped, washed and spun down, and resuspended in PBS. Samples were analyzed on a MACSQuant10 Flow Cytometry machine (Miltenyi). Gating strategy depicted in Fig. 7.

#### VSV-GFP Viral Supernatant Experiment

BMDMs were grown as described above in 150mm dishes. On day 8, following ~72 hours of puromycin treatment, media was removed and 10mL of PBS w/ 2mM EDTA was added. Cells were lightly scraped and 400,000 cells/well were replated in 12 well plates in BMDM media. Cells were left to adhere for 12 hours, before being infected at an MOI of 25 for 8 hours. Following infection, virus was removed and the cells were washed with PBS three times. Then, 500ul of BMDM media (DMEM, 20% FBS, 30% L929 condition media, and 1% Pen/Strep) was added to each well. 18 hours later, media was collected and stored at −80°C. To titer viral supernatant, Vero cells were plated in a 96-well plate at 30,000 cells per well in 90ul of D10 media. 12 hours after plating, 90ul supernatant was added to the 90ul of D10 at different dilutions. PFU/mL was calculated from a standard curve with a virus of known concentration.

#### Quantification and Statistical Analysis

All statistical analysis was performed in Python (version 2.7.9). Unless otherwise indicated in figure legends, statistical significance measurements were marked as follows: * denotes p < 0.05, ** denotes p < 0.01, *** denotes p < 0.001, and n.s. denotes not significant. RNA-Seq expression and splicing analysis as well as eCLIP analysis is described in more detail below.

#### RNA-Sequencing Analysis

Sequencing was performed on a HiSeq 2500 High Throughput Sequencer (Illumina). Single-end 50-mer reads were aligned using Tophat v2.1.1.^34^ Gene expression was determined using Cufflinks v2.2.1 and the FPKM (Fragments Per Kilobase Million) metric.^35^

#### Splicing Ratio and ∆SR Calculation

A custom script was written in Python using the HTSeq^36^ library to calculate Splicing Ratio. First, reads that map to an intron or exon feature are summed. To map to a feature, reads must have >1 bp overlap with the feature. If a read maps to more than one feature, such as in the case of a splice junction read, the read is split between the features. SR is calculated by taking the length normalized number of reads that map to each intron, divided by the average length normalized number of exon reads plus the length normalized intron value. When SR is equal to 0, this indicates a junction is completely spliced. In contrast, large SR values indicate intron retention. We use the SR value to calculate ∆SR, which is equal to SR(shBud13) - SR(Ctl). Values greater than 0 indicate the junction is more unspliced in the shBud13 sample, whereas values less than 0 indicate the junction is more unspliced in the Ctl sample. For the global analysis, in order for the ∆SR of a junction to be considered, it must pass through a number of filters. To account for transcripts that are annotated in Ensembl version 67, but not expressed, we set an FPKM threshold of 15. Further, a local normalized read count threshold on the upstream/downstream exons was implemented to ensure a level of sequencing depth needed to get accurate splicing values. To pass this threshold, the sum of the reads that map to the the upstream/downstream exons divided by the length of these exons must be ≥ 0.25.

#### ISG and Genome-Wide Analysis

ISGs used in Fig. 4 E-H were selected based on induction 2 hours after *in vivo* IFNα injection.^15^ We classified ISGs to be any gene with a fold change ≥ 3.5 following 2 hours of induction. Intron RPKM was calculated using a custom python script with the HTSeq library. In Fig. 5a, transcripts from the 30 min. TNFα data-set that had a junction with a ∆SR value above 0.15 were sorted into an ‘increased IR category (∆SR >0.15), whereas all other transcripts were sorted into an unaffected category (∆SR <-0.15). The selected data-set is representative of all time-points from the TNFα, Poly(I:C) and CpG datasets. A maximum entropy model was used to calculate 3’ and 5’ splice site strengths.^12^ To determine differences in 5’ splice site sequence for Bud13 dependent junctions, the nine base pair sequence near the 5’ splice site junctions for junctions that had a ∆SR >0.15 was compared to all expressed junctions (FPKM>1). The top Bud13 dependent junctions were plotted based on the average ∆SR value across all time-points from the TNFα data-set (Fig. 5D) as well as the Poly (I:C) data-set (Fig. SH). Junctions that had a ∆SR value <0.15 in a time-point were filtered out in the TNFα data-set, while junctions that had a ∆SR value <0.15 in two time-points were filtered out in the Poly (I:C) data-set. SVMBPfinder was used to determine BP related features (BP strength and distance from BP to 3’ splice site).^37^

#### eCLIP

Data for eCLIP experiments were downloaded from ENCODE Project Consortium.^17^ Analysis of eCLIP data is the same as has been described previously.^39^ Fold change of eCLIP read density compared to input read density along a normalized intron was calculated using ngs.plot.^40^ Peaks were called using CLIPper.^41^ Peaks were deemed significant if they were >3-fold enriched and had p-value<10-^5^. Peak locations were determined using a custom python script with the HTSeq library. Enriched GO terms were determined using Seten.^42^

#### Data and Software Availability

The next-generation sequencing data reported in this study will be deposited to the Gene Expression Omnibus (GEO). Upon completion of deposit, the accession number for this data will be provided.

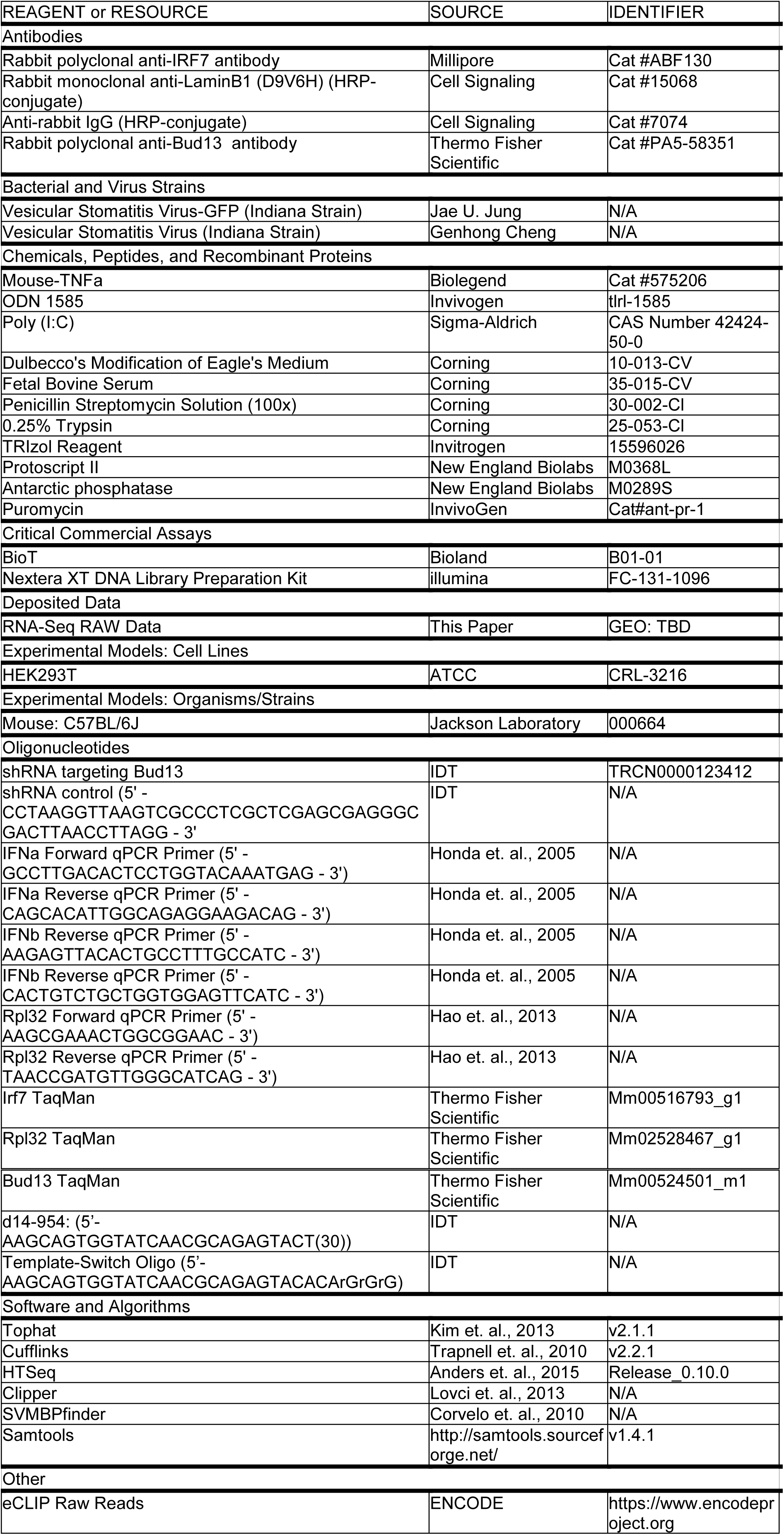

## SUPPLEMENTAL FIGURE LEGENDS

**Figure S1:**
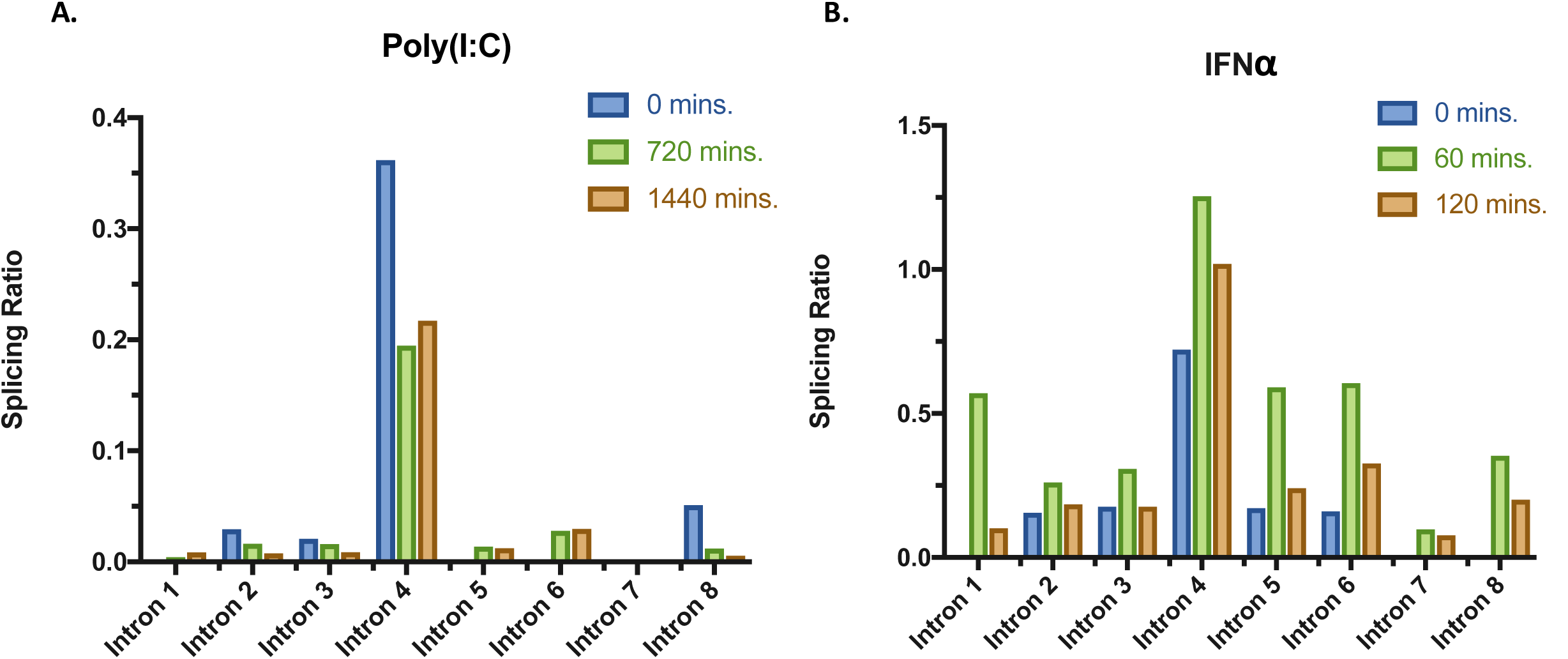
Splicing Ratios across all junctions in Irf7. Splicing ratios calculated for all junctions in the most abundant transcript of Irf7. Color represents time-point indicated in legend. **(A)** Poly(I:C) **(B)** IFNα.

**Figure S2:**
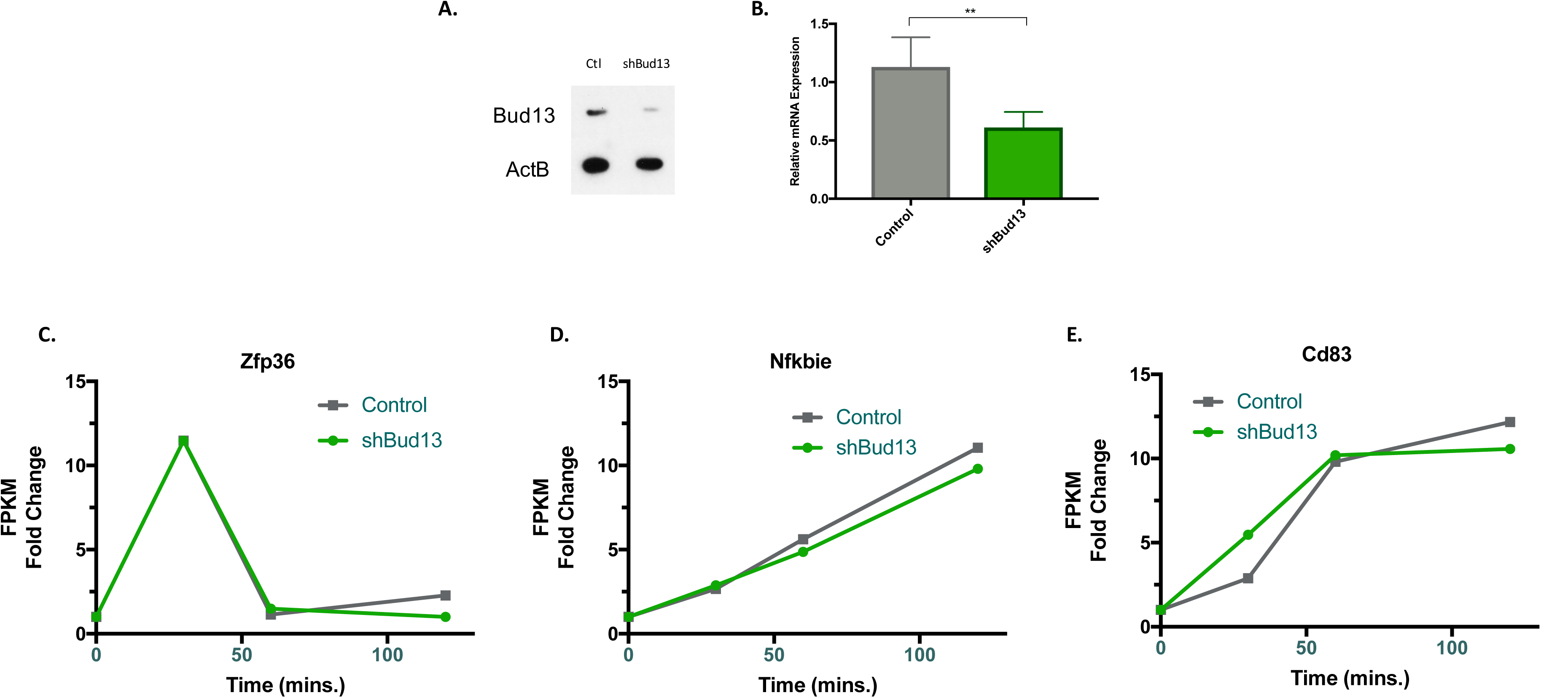
shBud13 knocks down Bud13 protein and mRNA. **(A)** Immunoblot analysis of Bud13 in BMDMs infected with control or shBud13. ActB serves as loading control. **(B)** qRT-PCR analysis of Bud13 mRNA in BMDMs infected with control or shBud13. **(C-E)** FPKM fold change with respect to time stimulated **(C)** Zfp36, **(D)** IκB?, and **(E)** CD83. shBud13 is shown in green, control is shown in grey. Data is representative of two individual experiments **(A, B)** and is shown as mean + SEM **(B)**. \**P* < 0.05, \*\**P* < 0.01 and \*\*\**P* < 0.001.

**Figure S3:**
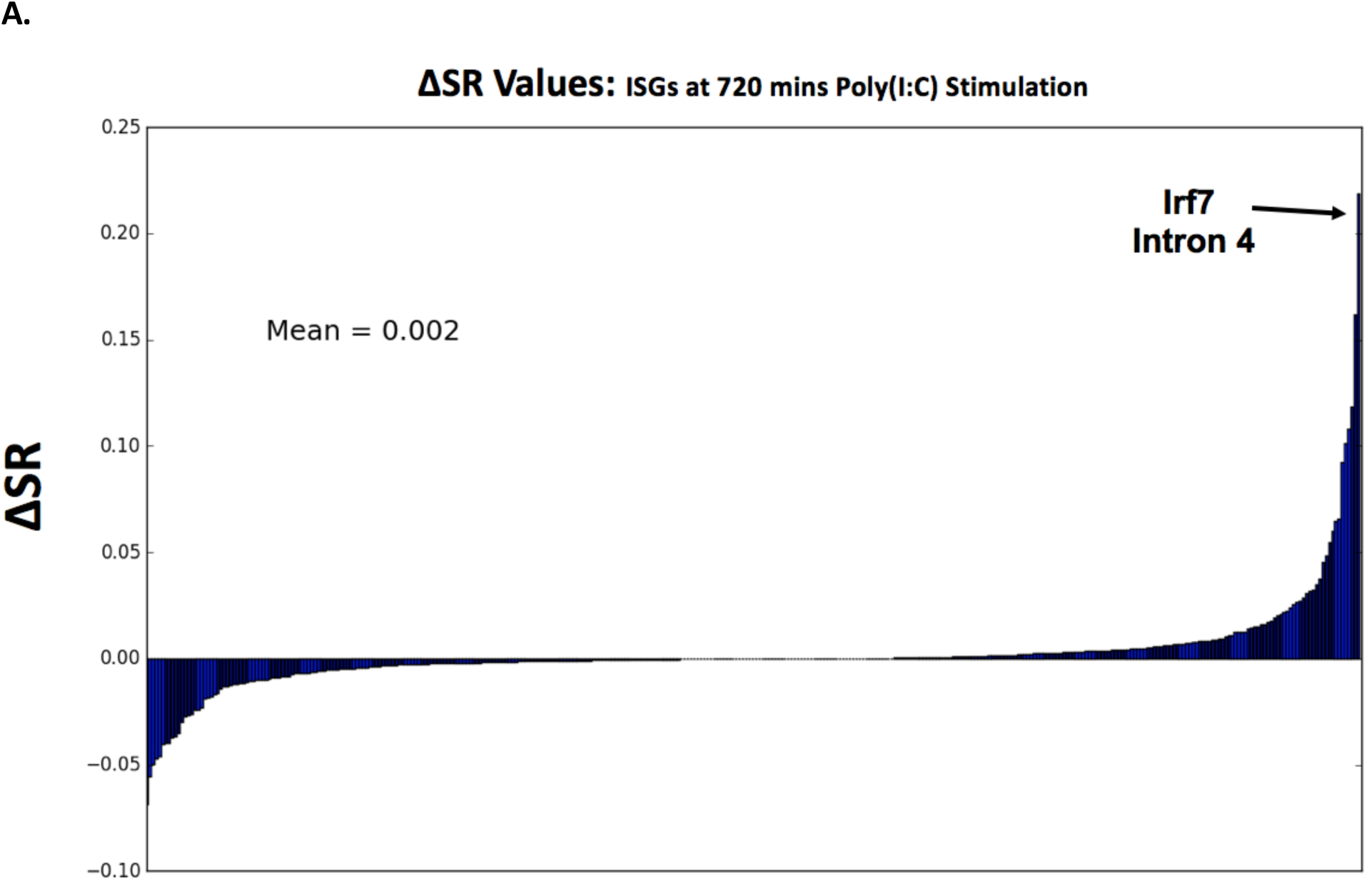
Irf7 Intron 4 is the most Bud13 knockdown affected junction of all ISGs. **(A)** ∆SR was calculated at 720 mins of poly(I:C) stimulation for each ISG junction that passed the transcript and local read count threshold (see methods). Mean ∆SR = 0.002, Median ∆SR = 0.

**Figure S4:**
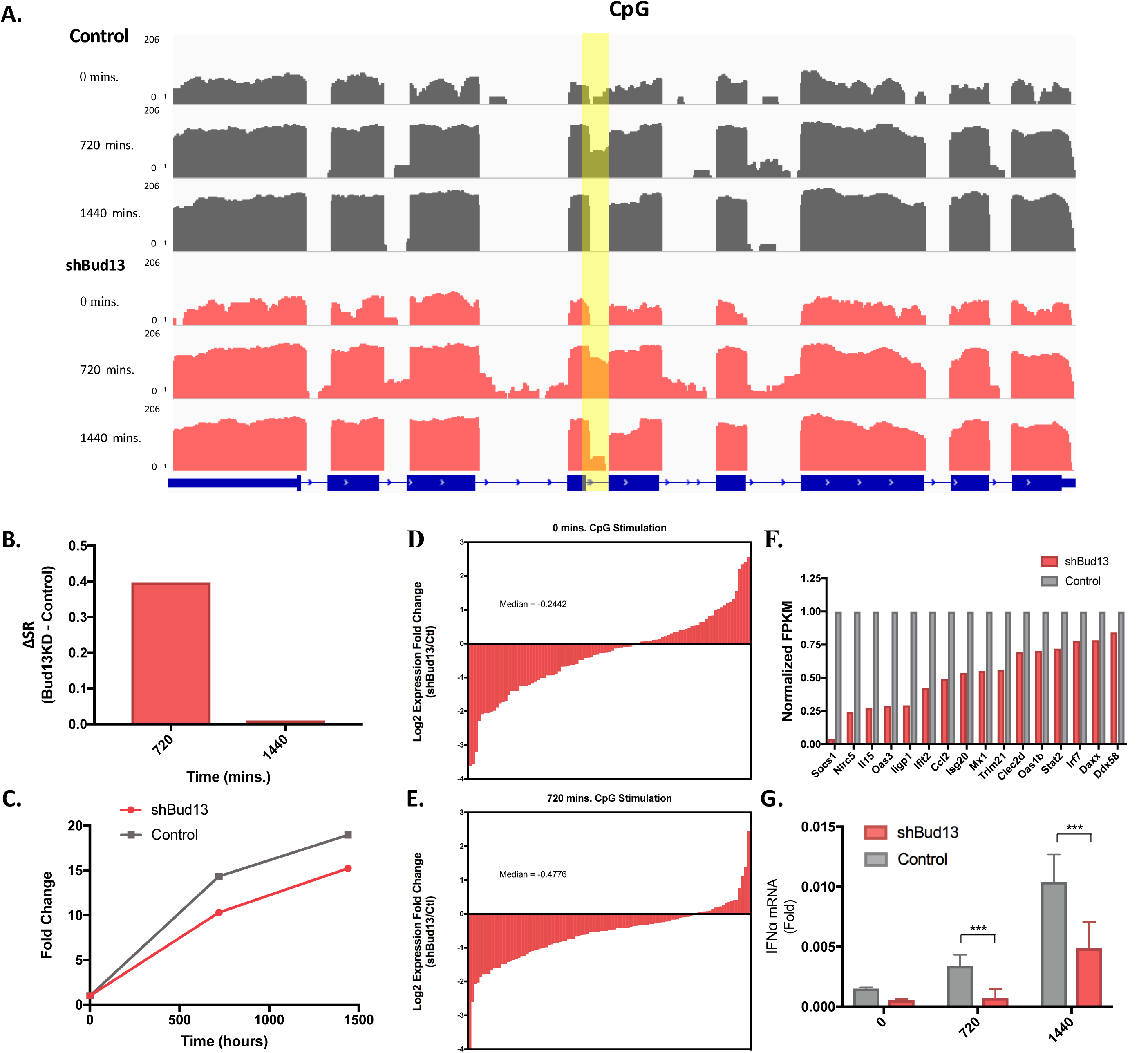
Bud13 knockdown alters the type I interferon response in response to CpG. **(A)** Histogram of mapped reads corresponding to the CpG-induced expression of Irf7. The poorly spliced fourth intron is highlighted. shBud13 samples are shown in pink. Control samples are shown in grey. **(B)** ∆SR calculated for the fourth intron of Irf7 at stimulated time-points (720, 1440 mins). **(C)** Irf7 FPKM fold change with respect to time stimulated. shBud13 is shown in pink, control is shown in grey. **(D)** Log_2_ expression fold change (shBud13/Control) for 119 ISGs (selected based on upregulation in response to IFNα) in unstimulated BMDMs (median = −0.2442). **(E)** As in **(D)** for stimulated BMDMs (720 mins CpG (median = −0.4776). Wilcoxon rank-sum between **(D)** and **(E)**, *P*< .001. **(F)** Normalized FPKM expression levels in shBud13 and control samples at 720 mins CpG stimulation for select ISGs. **(G)** RT-qPCR analysis of IFNα mRNA levels in unstimulated BMDMs and BMDMs stimulated with CpG for 720 and 1440 mins. Data is represented as mean + SEM. * denotes p < 0.05, ** denotes p < 0.01, and *** denotes p < 0.001 using a Student’s t test. Results are presented relative to those of Rpl32 **(G)**.

**Figure S5:**
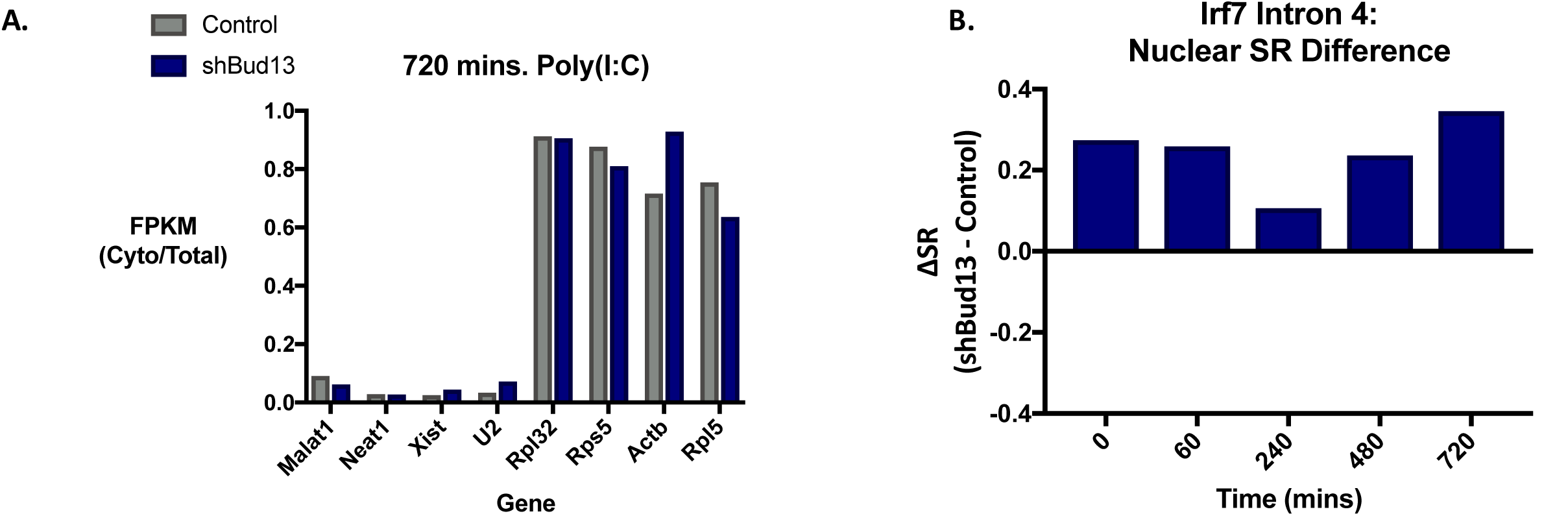
BMDM fractionation and effect of Bud13 on nuclear Irf7 splicing. **(A)** Ratio of cytoplasmic FPKM levels to cytoplasmic and nuclear FPKM levels for transcripts that are primarily nuclear (Malat1, Neat1, Xist, U2; left), and primarily cytoplasmic (Rpl32, Rps5, Actb, Rpl5; right) (BMDMs - 720 mins poly(I:C) stimulation). **(B)** ∆SR for intron 4 of Irf7 from nuclear fraction of unstimulated and poly(I:C) stimulated (15, 60, 240, 720 mins.) BMDMs.

**Figure S6:**
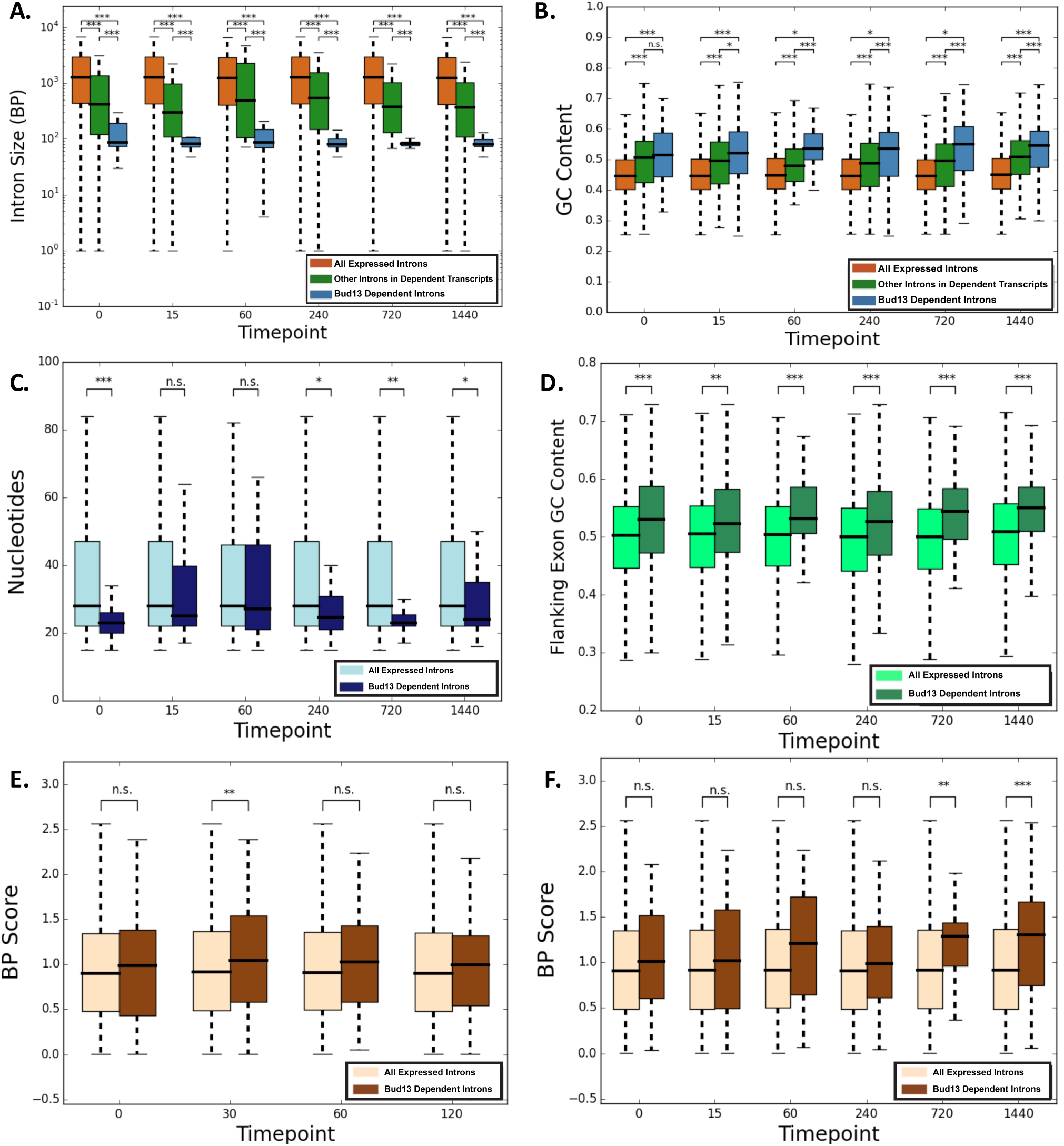
Supplemental global analysis of Bud13. **(A)** Size of intron for introns retained upon Bud13 knockdown (∆SR. > 0.15) (blue), in introns located in the same transcript as those affected by Bud13 (green), and in introns from all expressed transcripts (orange). **(B)** Same as **(A)** for GC content. **(C)** Flanking exon GC content for exons that flank introns retained upon Bud13 knockdown (∆SR. > 0.15) (dark green) as compared to exons that flank introns from all expressed transcripts (light green). **(D)** Distance from the branch point to the 3’ splice site for introns retained upon Bud13 knockdown (∆SR. > 0.15) (dark blue) as compared to introns from all expressed transcripts (light blue). **(A-D)** data from BMDM poly(I:C) stimulation. **(E)** Branch point score for introns retained upon Bud13 knockdown (∆SR. > 0.15) (beige) as compared to introns from all expressed transcripts (dark brown) in TNFα stimulated BMDMs. **(F)** Same as **(E)** but for poly(I:C) stimulated BMDMs. Box plots show median (center line), interquartile range (box) and tenth and ninetieth percentiles. \**P* < 0.05, \*\**P* < 0.01 and \*\*\**P* < 0.001 (Mann-Whitney *U*-test).

**Figure S7:**
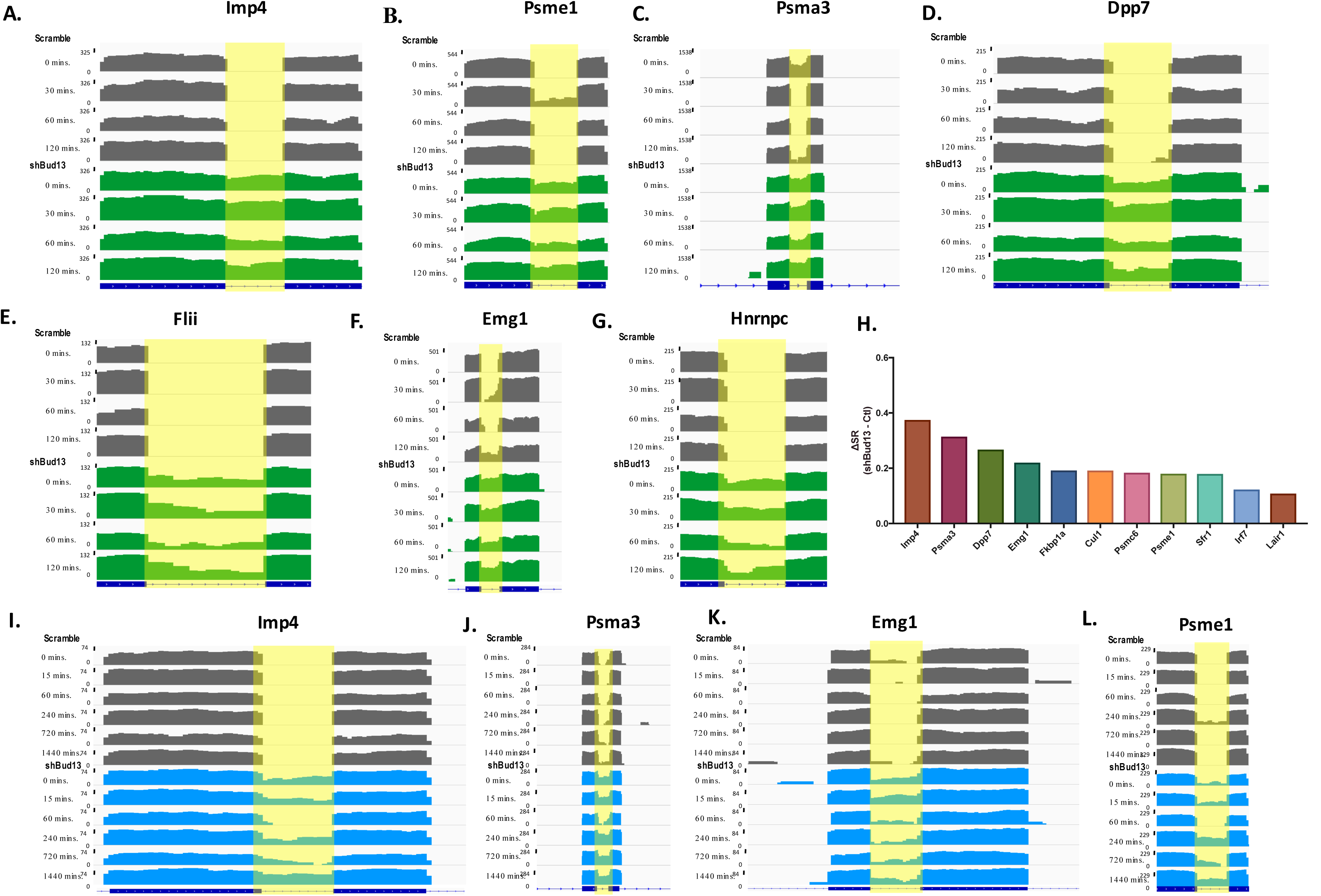
Histograms of mapped reads corresponding to hits identified in TNFα and PIC data-sets. **(A-G)** Histogram of mapped reads corresponding to the TNFα-induced expression Imp4, Psme1, Dpp7, Flii, Emg1, Hnrnpc, and Psma3 respectively. The poorly spliced intron in each transcript is highlighted. For all read density plots, reads are histogrammed in log_10_ scale and normalized to the maximum value across the stimulation. **(G)** Ranked bar chart showing genes with a junction most affected by Bud13 knock-down in all samples during PIC stimulation. **(I-L)** Histogram of mapped reads corresponding to the PIC-induced expression Imp4, Psma3, Emg1, and Psme1 respectively.

**Figure S8:**
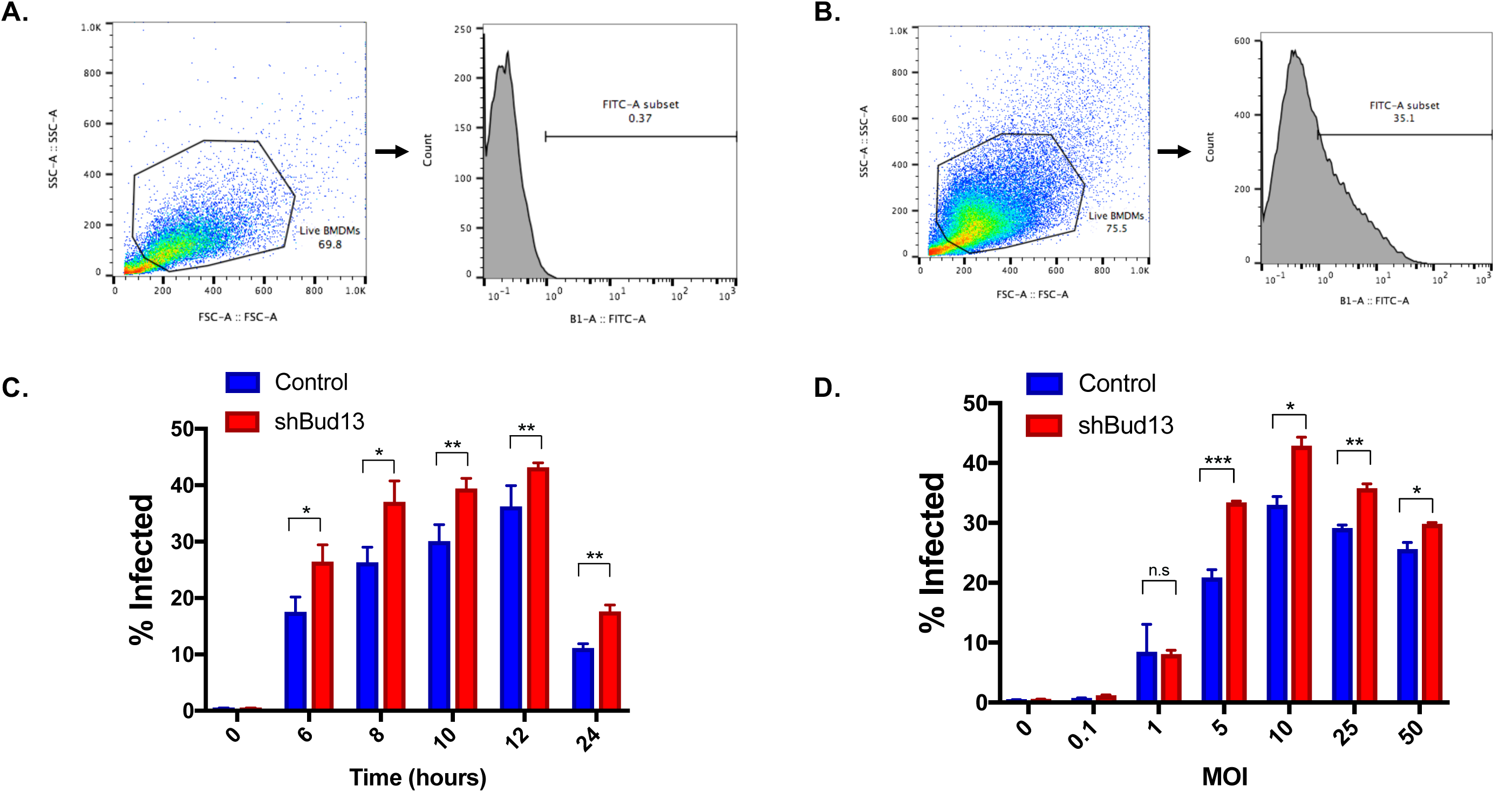
Bud13 knockdown alters the BMDM infection via VSV. **(A)** FSC/SSC plot showing the gating of live BMDMs in an uninfected control sample and the subsequent threshold used to calculate infectivity. **(B)** Same as in **(A)** but for a control sample infected with VSV-GFP for 12 hours. **(C)** Percent of live cells infected with VSV-GFP (MOI 10) in both control and shBud13 BMDMs across a 24-hour time-course. **(D)** Percent of live cells infected in both control and shBud13 BMDMs at 12 hours across a range of VSV-GFP MOIs.

